# Gene loss under constant cold reveals “natural knockout” loci in Antarctic notothenioid fishes

**DOI:** 10.64898/2026.06.06.730616

**Authors:** Vinita Lamba, Andrew J. Alverson, Jacob M Daane, Xuan Zhuang

## Abstract

**Background:** Gene loss is a major but often underappreciated mode of genome evolution. Genes loss can reflect relaxed selective constraint, neutral decay, compensation, or adaptive change; once fixed, however, they alter the inherited gene complement and can shape the evolutionary trajectories available to descendant lineages. Shared losses near the base of a radiation may therefore become part of the genomic background on which later specialization occurs. The Antarctic notothenioid (cryonotothenioid) radiation provides a powerful system for studying gene loss in an extreme polar environment: it diversified in the thermally stable Southern Ocean, and outgroups outside the clade allow ancestral gene presence to be inferred.

**Results:** Using whole-genome alignments and orthology-aware comparative analyses across 11 cryonotothenioids and four non-Antarctic notothenioid outgroups, we systematically identified gene-inactivating mutations and reconstructed their phylogenetic distribution. After applying stringent filters to reduce reference bias, mitigate assembly and annotation artifacts, and exclude loci with detectable transcriptomic support, we identified 30 high-confidence single-copy orthologs with loss of coding potential across sampled cryonotothenioids but intact coding sequences in outgroups. These clade-wide losses affect genes associated with lipid and amino acid metabolism, water transport, renal glucose reabsorption, skeletal mineralization, circadian regulation, and tRNA modification pathways. We also identified 12 additional single-copy orthologs lost across examined icefishes but retained in red-blooded notothenioids and outgroups, including genes associated with oxygen transport, iron handling, erythroid biology, and vesicular trafficking.

**Conclusions:** Our study provides a genome-wide catalog of coding-gene losses shared across sampled cryonotothenioids, together with additional lineage-specific losses in icefishes. Many of these losses occur in conserved genes associated with disease-relevant phenotypes in humans or model vertebrates, suggesting that Antarctic notothenioids offer a natural comparative system for studying how loss-of-function variants can persist in viable wild populations. These candidate natural knockouts provide testable hypotheses for how environmental buffering, paralog compensation, and pathway rewiring may shape vertebrate physiology under chronic cold.

## Background

Gene loss is a common feature of genome evolution that is frequently overlooked in studies of genome evolution [1–3]. Evolutionary novelty has traditionally been framed in terms of changes in protein sequence, gene regulation, or gene duplication, all of which can alter gene function or expression [4, 5]. Conversely, gene loss has often been interpreted primarily as a passive consequence of functional redundancy or relaxed selective constraint —a “use it or lose it” view. For example, the loss of intestinal fat digestion genes such as *MOGAT2* and *FABP6* in fruit bats is likely a consequence of relaxed selection following specialization on a fat-poor frugivorous diet [6]. A complementary “less is more” perspective emphasizes that gene loss can also contribute directly to phenotypic evolution by removing functions that are costly or disadvantageous in a particular ecological context [7]. For example, the loss of *AMPD3* in sperm whales may increase erythrocyte ATP and facilitate oxygen unloading during deep dives, whereas loss of *MMP12* in fully aquatic mammals may preserve lung elastin and support rapid exhalation at the surface [6]. Similarly, gene loss has served as a gateway to rapid adaptation and evolutionary innovation in yeast [8]. Together, these views highlight gene loss as a fundamental component of genome evolution, with the potential to shape lineage trajectories through multiple, non-mutually exclusive ways.

From a molecular evolutionary perspective, gene retention and gene loss largely reflect differences in selective constraint. Genes with essential or broadly required functions are typically maintained by purifying selection and therefore show high sequence conservation [9, 10]. In contrast, genes with reduced, redundant, or context-dependent functions can tolerate more sequence and structural variation, making them more likely to accumulate loss-of-function mutations and eventually undergo pseudogenization or deletion [11, 12]. Importantly, the evolutionary consequences of losing a gene are not limited to that gene alone: because genes function within interconnected molecular and cellular networks, the loss of a single component can have system-level effects and reshape the constraints and opportunities available to descendant lineages [8]. Thus, gene loss need not always be interpreted as a direct adaptive response; once fixed, losses become part of the inherited genomic starting point for subsequent evolution. Accordingly, genome-wide surveys of gene loss can reveal how disruptions accumulate across interacting pathways, providing a systems-level complement to gene-by-gene or single-trait studies.

Antarctic notothenioid fish offer a powerful system for investigating gene loss during adaptive radiation in an extreme but unusually stable environment. The Antarctic clade diverged during the Early Miocene; recent time-calibrated phylogenies derived from genome-wide sequence data estimate that their rapid adaptive radiation began much later, at approximately 10.7 million years ago (MYA) [13] following long-term Southern Ocean cooling and thermal isolation associated with the Antarctic Circumpolar Current (ACC) [14, 15]. The freezing of the Southern Ocean is thought to have reduced much of the pre-existing fish fauna, creating ecological opportunity for notothenioids. Following extensive physiological and ecological specialization, the Antarctic clade (Cryonotothenioidea) today comprises over 120 species, serves keystone ecological roles, and dominates the region’s fish biomass [16].

The Southern Ocean’s chronic cold and long-term thermal stability shaped notothenioid genomes at multiple levels. Cryonotothenioids have substantial variation in genome size, driven by clade-wide bursts of multiple transposable element (TE) families [13]. Furthermore, their evolutionary trajectory is marked by chromosomal rearrangement, karyotype evolution, and other large-scale genomic changes [17, 18]. A hallmark example of adaptation is the evolution of antifreeze glycoprotein (AFGP) genes, representing a genomic innovation that enabled survival in subzero waters and opened the path to notothenioid diversification [19]. In contrast, loss of function such as the inducible heat shock response (HSR) [20–22], erosion of circadian gene repertoires [23, 24], and hemoglobins in the icefishes lineage [25–28] reflects specialization to a permissive, oxygen-rich Antarctic environment, where functions essential in other vertebrates became dispensable. Despite these prominent examples, coding-gene inactivation in Cryonotothenioidea remains incompletely characterized beyond a handful of well-studied loci and trait-focused analyses [29, 30].

In this study we performed whole-genome alignments and orthology-aware comparative analyses to systematically identify gene-inactivating mutations across 11 Antarctic and four non-Antarctic notothenioid species. Leveraging these non-Antarctic taxa as phylogenetic anchors enables robust inference of Antarctic clade–specific inactivation events. By reconstructing the phylogenetic distribution of disruptions and applying stringent filters to minimize reference bias and assembly/annotation artifacts, we identify gene losses shared across Cryonotothenioidea as well as additional losses restricted to the highly derived icefish lineage. Together, this genome-wide analysis establishes Antarctic notothenioids as a comparative system for studying gene loss under constant cold and for prioritizing candidates for future functional testing.

## MATERIALS AND METHODS

### Genome and transcriptome resources

To systematically identify genome-wide gene inactivation across Antarctic notothenioids, we retrieved whole-genome assemblies for 11 sampled members of the cryonotothenioid clade (*Dissostichus eleginoides*, *Dissostichus mawsoni*, *Lepidonotothen nudifrons*, *Trematomus bernacchii*, *Pogonophryne albipinna*, *Harpagifer antarcticus*, *Gymnodraco acuticeps*, *Champsocephalus esox*, *Champsocephalus gunnari*, *Chaenocephalus aceratus*, and *Pseudochaenichthys georgianus*) and four taxa outside the cryonotothenioid clade (*Cottoperca gobio* [NCBI assembly name; synonym of *Cottoperca trigloides*] [31, 32], *Bovichtus diacanthus*, *Bovichtus variegatus*, and *Eleginops maclovinus*) from the NCBI GenBank and RefSeq databases (**Table S1**). Here, we use “cryonotothenioid clade” in a phylogenetic sense to refer to the Antarctic notothenioid radiation, rather than as a strict statement of present-day geographic distribution for every sampled species. These assemblies were strategically selected to represent all five Antarctic families together with available non-Antarctic notothenioid outgroups, providing a phylogenetic framework for reconstructing the ancestral gene complement at the base of Cryonotothenioidea. This design allowed us to identify genes retained outside the cryonotothenioid clade but lost across sampled cryonotothenioids, while also distinguishing additional losses restricted to the derived icefish lineage. Assembly quality was evaluated using BUSCO (v5.4.2) [33], and all selected genomes recovered more than 93% of conserved single-copy orthologs, supporting their suitability for comparative, synteny-based analyses.

To support transcript-level validation of candidate gene losses, we used publicly available RNA-seq datasets representing Antarctic notothenioids and a closely related non-Antarctic species from the NCBI Sequence Read Archive (SRA) (**Table S2**). Collectively, these datasets span multiple tissues (liver, heart, brain, gill, spleen, and muscle), developmental stages (embryonic, juvenile, and adult), and both sexes, thereby broadening the range of expression contexts available for evaluating whether putatively inactivated genes retain detectable transcription.

### RNA-seq processing and transcriptomic assembly

For datasets requiring processing from raw reads, sequencing reads were downloaded using SRA Toolkit (v3.0.0) (https://github.com/ncbi/sra-tools). These included raw reads from the Antarctic icefishes *Champsocephalus gunnari* and *Champsocephalus esox* (PRJNA859929), as well as the non-Antarctic species *Eleginops maclovinus* (SRX2523921). Reads were subjected to adapter removal and quality trimming using Trimmomatic (v0.39) [34], with the parameters ILLUMINACLIP:TruSeq3-PE.fa:2:30:10 LEADING:3 TRAILING:3 SLIDINGWINDOW:4:15 MINLEN:36. The retained high-quality reads were assembled de novo into transcript contigs using Trinity (v2.15.1) [35]. In parallel, high-quality reads were aligned to their respective reference genomes using the splice-aware aligner STAR [36] to assess exon-level coverage and splice-junction support across candidate loci.

The resulting transcript assemblies and RNA-seq alignments were used as an independent validation layer for TOGA-inferred gene losses. Absence of transcriptomic support was interpreted conservatively as supportive, but not definitive, evidence of gene non-functionality, because expression may depend on tissue, developmental stage, sex, or environmental condition.

### Species Phylogeny Reconstruction

To provide the phylogenetic framework used to interpret losses shared across Cryonotothenioidea versus additional losses restricted to icefishes, we inferred species relationships from the predicted proteomes of all 15 taxa. Protein sequences were clustered into orthogroups with OrthoFinder [37] and OrthoFinder was run in MSA mode (-M msa -A muscle -T iqtree). Multiple sequence alignment was performed using MUSCLE [38], and gene trees were inferred with IQ-TREE [39]; the species tree was estimated using the STAG method implemented in OrthoFinder [40], based on 12,374 orthologous groups. The resulting tree topology was congruent with recent phylogenomic studies of notothenioid fishes [14, 41].

### Whole-genome alignment and orthology inference

To identify lost genes across cryonotothenioids while minimizing false inferences due to annotation or assembly errors, we used a whole-genome alignment based comparative genomics framework that integrates sequence similarity and conserved synteny. We define “outgroups” here as non-Antarctic notothenioids outside Cryonotothenioidea that are used to infer ancestral gene presence and to polarize cryonotothenioid clade–specific loss events. We analyzed genome assemblies from 11 cryonotothenioid species together with four non-cryonotothenioid species as outgroups (**Table S1**). Two of these outgroup taxa, *Cottoperca gobio* (GCA_900634415.1) and *Eleginops maclovinus* (GCF_036324505.1), also served as reference genomes for TOGA projection; the remaining non-Antarctic taxa (e.g., Bovichtus spp.) were included as independent comparisons to reduce reference bias and to guard against lineage-specific loss in any single non-Antarctic lineage. Each query assembly was aligned independently to each reference genome using the make_lastz_chains pipeline [42, 43] (https://github.com/hillerlab/make_lastz_chains), which produces high-quality pairwise alignment chains suitable for orthology inference from genome alignments.

Orthologous loci and gene-inactivating mutations were inferred using TOGA (Tool to infer Orthologs from Genome Alignments) [44] (https://github.com/hillerlab/TOGA). TOGA uses the reference gene annotation together with the reference–query alignment chains to locate the orthologous region in each query genome and to project reference transcript models onto that locus. TOGA then evaluates gene integrity across annotated transcript isoforms, focusing on the middle 80% of the coding sequence (CDS). Based on the projected gene structure and the presence of predicted gene-inactivating mutations, TOGA assigns each gene to predefined status classes, including intact (I), partially intact (PI), lost (L), or uncertain loss (UL). TOGA designates genes as lost (L) when multiple inactivating mutations (e.g., frameshift insertions or deletions, premature stop codons, exon deletions, or splice-site disruptions) were detected within the middle 80% CDS region. Genes with a single inactivating mutation in this region were classified by TOGA as uncertain losses (UL), reflecting a lower confidence call that could also arise from local assembly or projection artifacts.

TOGA also reports categories that reflect incomplete sequence recovery rather than clear gene loss. A locus is classified as partially intact (PI) when part of the CDS is missing but no inactivating mutation is detected within the internal 80% of the aligned CDS. It is classified as missing (M) when a large fraction of the predicted CDS is absent or masked (typically due to assembly gaps), and as partially missing (PM) when much of the projected locus extends beyond scaffold boundaries. For downstream analyses, we treated PI calls as not lost unless additional evidence supported inactivation.

### Mitigating reference bias and defining high-confidence gene losses

To minimize false-positive loss calls, we applied additional stringency filters (**Figure 1**).

**Figure 1.**
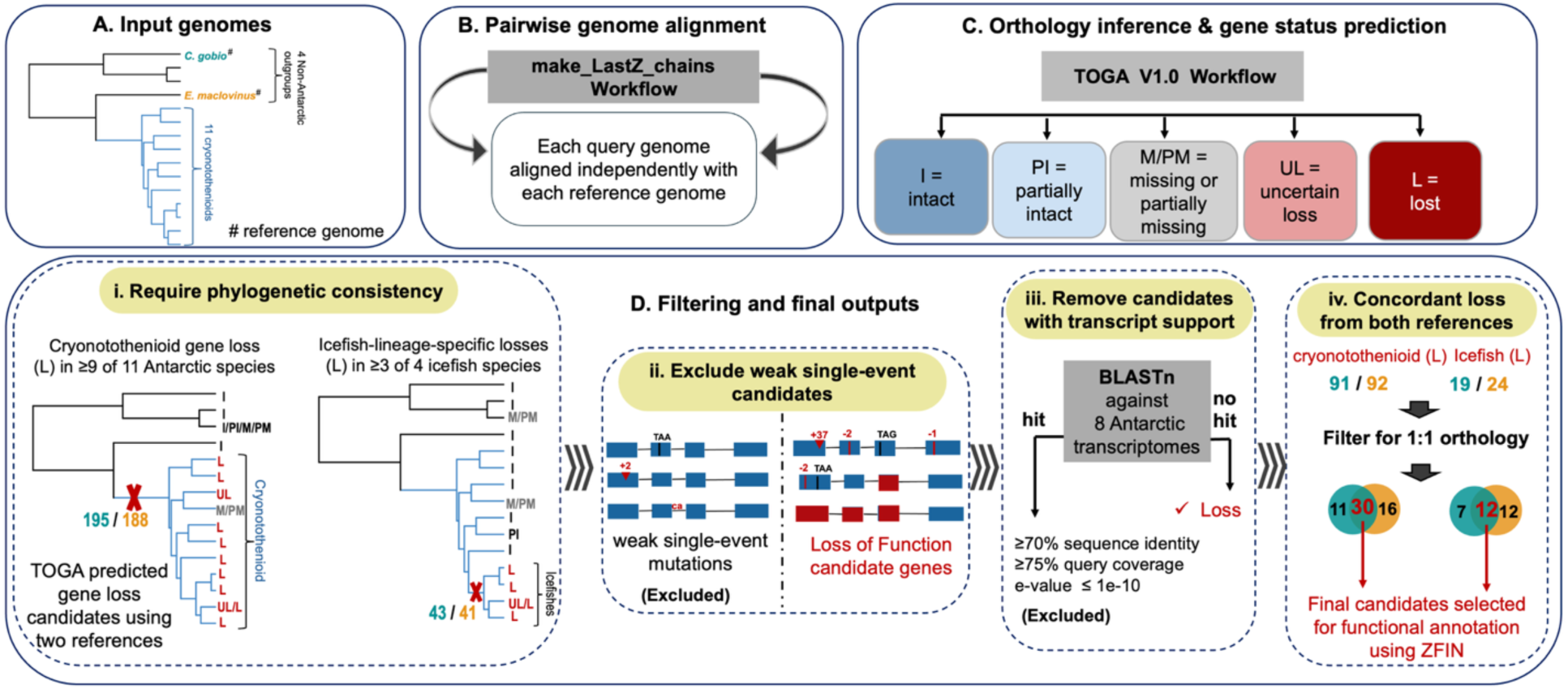
Workflow for inferring high-confidence gene losses in cryonotothenioids. **A)** Input genome sampling included 11 cryonotothenioids and four outgroups outside the cryonotothenioid clade. *Cottoperca gobio* and *Eleginops maclovinus* were used both as outgroups and as independent reference genomes for downstream analyses, indicated by #. **B)** Each query genome was aligned independently to each reference genome using the make_lastz_chains workflow to generate pairwise whole-genome alignment chains. **C)** TOGA was used for orthology inference and gene-status prediction from the reference–query genome alignments. Gene-status classes include intact (I), partially intact (PI), missing or partially missing (M/PM), uncertain loss (UL), and lost (L). **D)** Candidate losses were filtered through four sequential steps. First, phylogenetic consistency was required: candidate cryonotothenioid clade-wide losses were required to be classified as lost in at least 9 of 11 cryonotothenioids, while candidate icefish-lineage-specific losses were required to be classified as lost in at least 3 of 4 examined icefishes and retained in red-blooded cryonotothenioids and outgroups. Second, weak candidates supported only by isolated single-event mutations were excluded. Third, candidates with transcriptomic matches in eight Antarctic transcriptomes were removed using BLASTN thresholds of ≥70% sequence identity, ≥75% query coverage, and E-value ≤1e−10. Finally, candidates were required to show concordant loss calls across both reference-based analyses and were filtered for 1:1 orthology. Venn diagrams summarize the overlap between the two reference-based analyses, with the central intersections representing the final candidate sets selected for functional annotation.

First, candidate cryonotothenioid clade–wide losses were required to be classified as lost (L) in at least nine of the eleven species in the cryonotothenioid clade, while allowing up to two species to be classified as uncertain loss (UL), partially intact (PI), or missing (M). This allowance accounts for variation in assembly continuity, because several genomes are scaffold-level rather than chromosome-level assemblies. Candidate clade-wide losses were also required to retain intact coding potential in non-cryonotothenioid outgroups. For icefish-lineage-specific losses, we applied analogous criteria, requiring candidate genes to be classified as lost in at least three of the four examined icefish species while remaining intact in red-blooded cryonotothenioids and non-cryonotothenioid outgroups.

Second, we removed borderline candidates in which loss status was supported only by a small number of predicted inactivating mutations, such as an isolated frameshift, premature stop codon, or limited splice-site changes. Such patterns may reflect sequencing error, local misassembly, or annotation/projection artifacts rather than true loss of coding potential.

Third, we used transcriptomic evidence to remove candidates with evidence of continued transcription in Antarctic species. To validate inferred losses at the transcript level, all candidate genes were queried against eight available Antarctic transcriptomes spanning four notothenioid families (**Table S2**) using BLASTN [45]. Candidates were excluded if they produced transcriptomic matches meeting ≥70% sequence identity, ≥75% query coverage, and E-value ≤1e-10. Candidates lacking such matches were retained as putative non-functional candidates.

Finally, because TOGA-based gene status inference can depend on the choice of reference genome, we performed the full analysis independently using *Cottoperca gobio* and *Eleginops maclovinus* as references. TOGA predictions generated from each reference were compared across all species, and only candidates satisfying the relevant loss criteria in both reference-based analyses were retained as high-confidence losses. Detailed per-gene status calls and filtering outcomes are provided in Tables S4–S7.

Together, this multi-layered filtering strategy was used to define high-confidence cryonotothenioid clade-wide and icefish-lineage-specific gene losses supported by phylogenetic consistency, sufficient coding-sequence disruption, absence of detectable transcriptomic support, and concordant inference across both reference genomes.

### Functional annotation and phenotypic inference

Candidate genes were retained only if they were consistently identified as lost across both reference genomes and exhibited no hits in any of the available the transcriptome data. For these validated genes, we utilized g:Orth module from g:Profiler web server [46] (http://biit.cs.ut.ee/gprofiler/orth) to determine their 1:1 orthology status in *Danio rerio* and *Homo sapiens*. To determine the biological roles of the 30 inactivated single-copy orthologs, we performed a manual curation using functional databases and primary literature. Specifically, we used the Zebrafish Information Network (https://zfin.org) and primary literature to identify previously determined biological functions and or documented knockout phenotypes in *Danio rerio* and other model vertebrates. This comparative database approach was used to generate hypotheses about the potential physiological and phenotypic consequences associated with the loss of these candidate genes in Antarctic notothenioid.

## Results

### Systematic detection of gene losses in Antarctic notothenioids

We analyzed 11 cryonotothenioid genomes and four non-Antarctic relatives using an orthology-aware, synteny-guided comparative framework (TOGA pipeline) [44] to characterize genome-wide patterns of gene loss across the clade. Protein-coding genes were annotated using two non-Antarctic reference genomes—*Cottoperca gobio* (GCA_900634415.1; 21,662 genes) and *Eleginops maclovinus* (GCF_036324505.1; 22,649 genes)—which represent outgroups outside cryonotothenioids. These outgroups were used to infer the ancestral gene complement at the base of the Antarctic radiation and to polarize Antarctic clade–specific gene losses. We used a multi-step filtering strategy to infer gene losses shared across Cryonotothenioidea, as well as additional lineage-specific losses within Antarctic icefishes, the white blooded notothenioids (Channichthyidae).

Each gene in each species was characterized as intact (I), partially intact (PI), lost (L), uncertain loss (UL), or missing (M). Across the Antarctic radiation, the largest gene-status category was intact, with ≥67% classified as intact using *Cottoperca gobio* as the reference and >75% using *Eleginops maclovinus* as the reference. *D. mawsoni* was an exception, with fewer than 40% of genes assigned intact coding status; this pattern was driven largely by an elevated uncertain-loss category and was not interpreted as evidence of extensive true gene loss. Smaller fractions of genes across the clade were classified as lost (5–7%), missing (2–5%), and uncertain loss (20–25%), with broadly similar distributions across species and reference annotations (**Figure 2B**). Among all gene status categories, *L. nudifrons* showed the highest number of missing (M) genes, consistent with a larger fraction of loci falling in assembly gaps or poorly resolved regions (**Figure 2B**).

**Figure 2.**
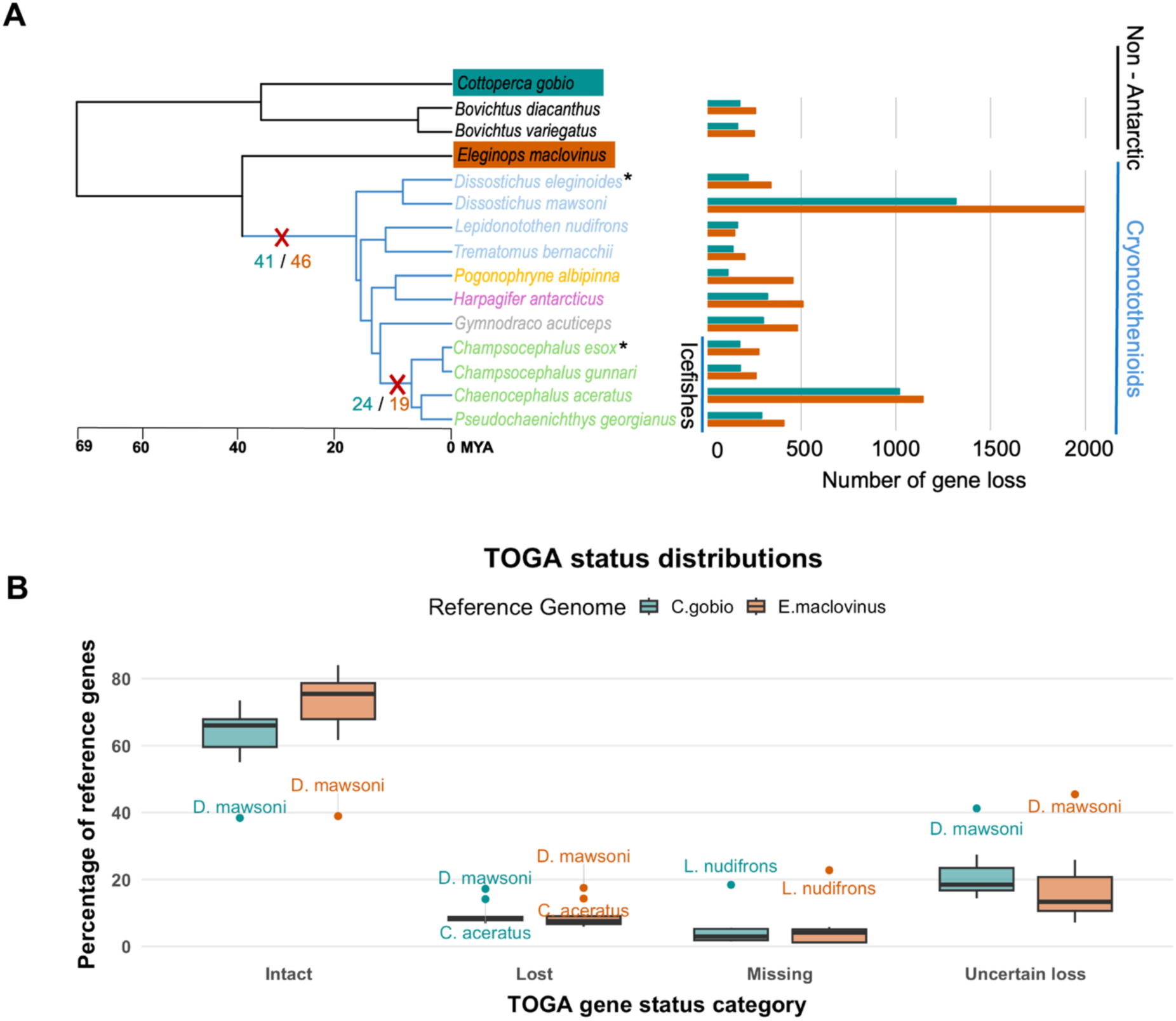
Genome-wide annotation and classification of gene status across Antarctic notothenioids. **A)**. A phylogeny of notothenioids, including four non-Antarctic outgroup species and representatives of the five cryonotothenioid families. The topology was reconstructed from single-copy orthologs inferred by OrthoFinder, and divergence times were added using TimeTree. Highlighted outgroups indicate the two reference genomes used for TOGA projection. The bar plot shows the number of genes classified as lost in each species using each reference genome. Red crosses mark candidate loss inferred on the branch leading to the sampled cryonotothenioids and on the icefish lineage; numbers beside the crosses indicate the corresponding loss counts from the two reference-based analyses. **B)**. Status of 21,662 protein-coding genes from *Cottoperca gobio* and 22,649 protein-coding genes from *Eleginops maclovinus* inferred by TOGA across the sampled cryonotothenioid assemblies. For each species, genes were classified as intact (I; clearly intact), partially intact (PI; some fraction of CDS is missing, but the gene is likely intact), lost (L; containing multiple gene-inactivating mutations such as premature stop codons, frameshifts, splice-site disruptions, or exon deletions), uncertain loss (UL; single inactivating mutation, indicating lower-confidence loss calls), and missing (M; substantial CDS missing or masked, often reflecting assembly gaps). The x-axis shows TOGA gene-status categories, and the y-axis shows the percentage of reference genes assigned to each category. Points indicate species-level values; labels identify notable outliers. Color corresponds to reference species. Five cryonotothenioid families: Nototheniidae (blue), Artedidraconidae (orange), Harpagiferidae (pink), Bathydraconidae (grey) and Channichthyidae (green). The * symbol indicates sampled members of the cryonotothenioid clade with secondarily temperate or sub-Antarctic distributions.

Using the high-confidence filtering strategy described in Methods, we identified 30 genes inferred to have lost coding potential across the Antarctic clade while retaining intact coding potential in all non-Antarctic outgroups (**Figure 2A**; **Figure S5, Table S4-S5**). These candidates were supported by concordant loss calls from both reference-based analyses, broad phylogenetic consistency across Antarctic species, absence of strong transcriptomic support, and extensive coding disruption. Within the Antarctic radiation, we also identified 12 single-copy orthologs consistently predicted as lost across all examined icefishes but retained in red-blooded notothenioids and non-Antarctic outgroups. These lineage-specific gene losses are characterized by multi-exon deletions and/or multiple frameshift mutations, suggesting inactivation at the ancestral node of the icefish family (**Figure 2A**; **Figure S6, Table S6-S7**).

### Functional annotation of ancestral and lineage specific gene losses and candidate gene selection

To explore the potential biological context of ancestral gene losses, we annotated the final set of 30 ancestral losses using Gene Ontology databases (ShinyGo 0.80) and manual curation. Although no standard Gene Ontology (GO) categories were significantly enriched after multiple-testing correction (FDR < 0.05), many of the lost genes have known functions in vertebrate physiological systems relevant to Antarctic notothenioid biology, including lipid metabolism, proline catabolism, renal glucose handling, skeletal mineralization, circadian regulation, and tRNA modification. Several of these genes are also associated with human disease phenotypes, including metabolic dysfunction, hyperprolinemia, renal glycosuria, skeletal disorders, and circadian rhythm disorders (**Table 1**). Icefish-specific losses included genes associated with oxygen transport, erythropoiesis, vesicular trafficking, and thrombocyte function, with corresponding human disease associations including thalassemia, congenital dyserythropoietic anemia, and Bernard-Soulier syndrome (**Table 1, Table S3**). We used these functionally characterized ancestral and icefish-specific losses to select representative candidate “natural knockouts” for detailed analysis below.

**Table 1.**
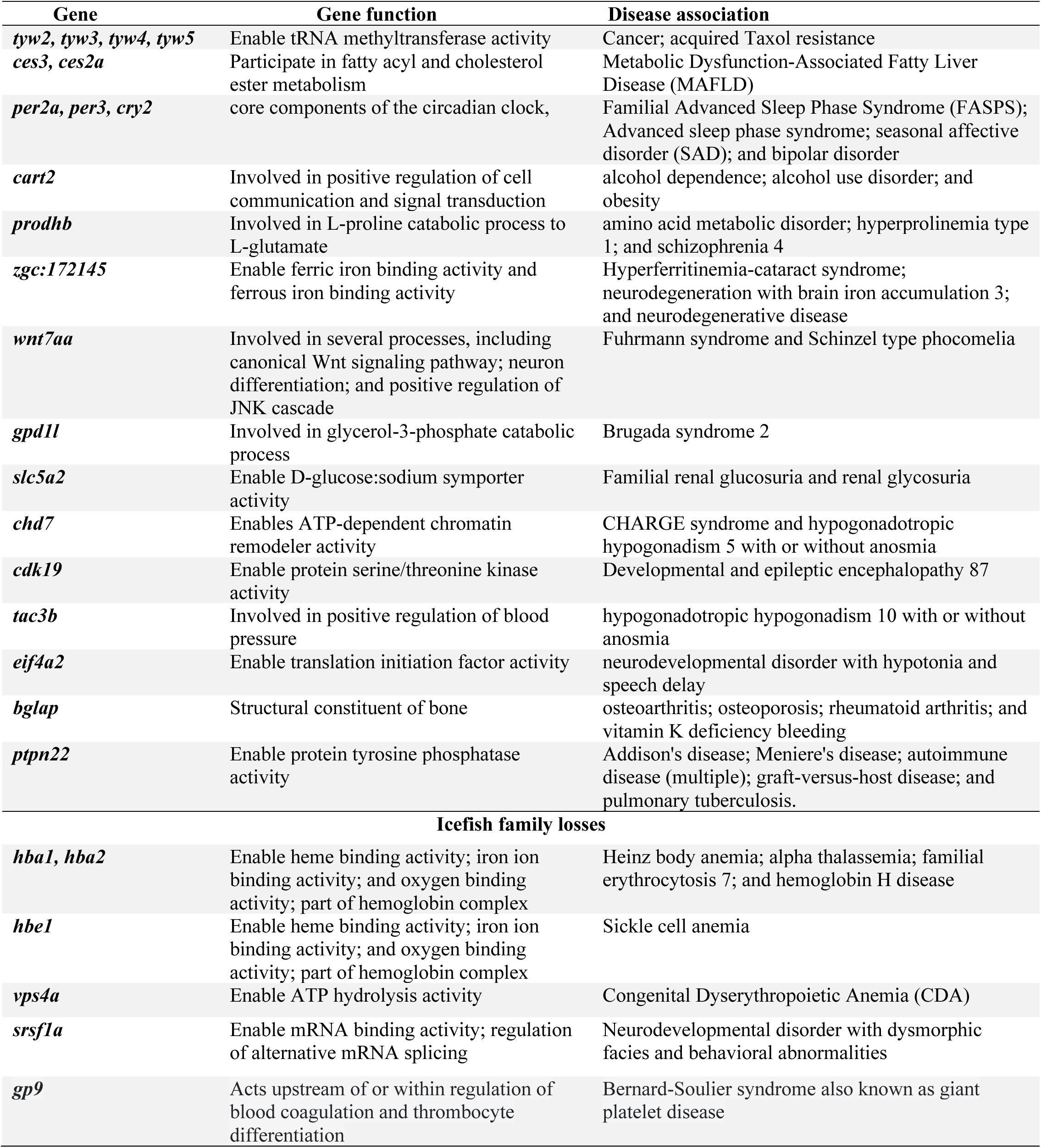
Summary of gene function and the associated disease in human.

### Genomic disruption of hepatic lipases (*ces3* and *ces2a*) involved in triacylglycerol regulation

A notable feature of Antarctic notothenioid lipid biology is the wide range of total lipid content across species and ecological niches—ranging from massive accumulations in pelagic species like *Aethotaxis mitopteryx* (61.4% of dry mass) to significantly lower stores in benthic lineages like *Dolloidraco longedorsalis* (14.5%). While total lipid mass varies significantly among notothenioids which strongly correlates with their distinct ecological niches, triacylglycerols (TAGs) consistently emerge as the dominant lipid class across the clade [47–49].

The *ces3* gene encodes a triacylglycerol hydrolase (TGH) and the *ces2* gene encode hepatic carboxylesterases; both are involved in neutral lipid hydrolysis and lipid homeostasis [50–52]. We identified the inactivation of both *ces3* and *ces2a* in cryonotothenioids, while both genes remain fully intact in all four non-Antarctic outgroups. For *ces3*, a shared premature stop codon in exon 8 suggests an ancestral inactivating mutation, followed by additional lineage-specific frameshift mutations (**Figures 3A, 3B**). Similarly, the 11-exon *ces2a* contains multiple frameshift mutations and premature stop codons; a shared 4-nucleotide insertion (‘CTGT’) in exon 6 supports an ancestral gene-inactivating mutation predating diversification of the Antarctic clade (**Figure 3C**).

**Figure 3.**
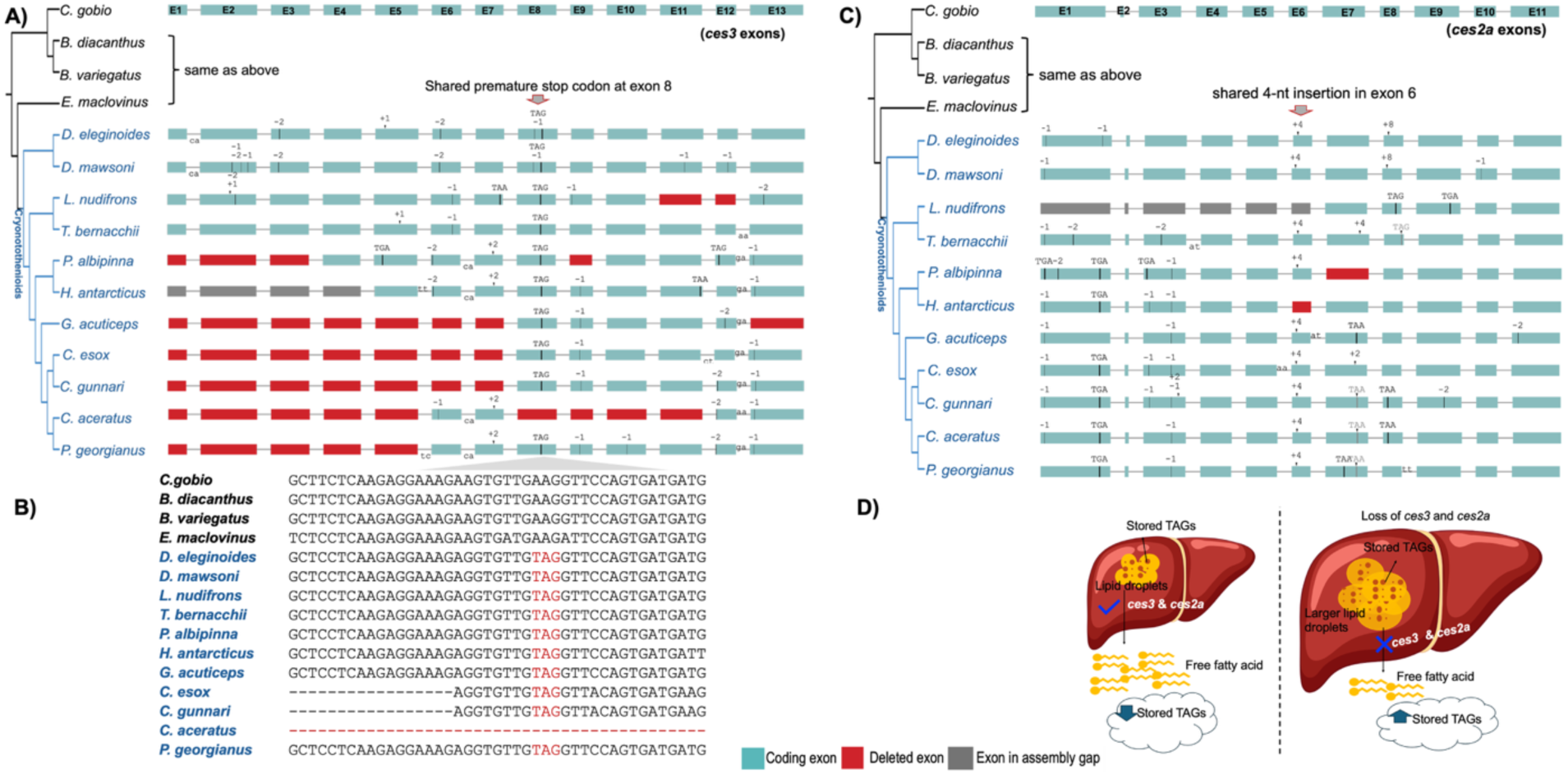
Loss of hepatic carboxylesterases and altered triacylglycerol turnover. **A)** *ces3* gene model comparison across outgroups and cryonotothenioids. Teal boxes indicate retained coding exons, red boxes indicate deleted exons, and grey boxes indicate missing or assembly-gap regions. A premature stop codon in exon 8 is shared across sampled cryonotothenioids, followed by additional lineage-specific frameshifts, splice-site changes, or exon deletions. **B)** Multiple sequence alignment of *ces3* exon 8 highlighting the shared premature stop codon (TAG) in sampled cryonotothenioids. The dashed red line indicates exon deletion in *C. aceratus*. **C)** *ces2a* gene model comparison showing a shared 4-nucleotide insertion in exon 6 across cryonotothenioids, together with additional coding-sequence disruptions. **D)** Conceptual model illustrating how loss of CES-mediated TAG hydrolysis may alter hepatic lipid turnover and provide a permissive metabolic context for TAG-rich lipid storage in lineages where lipid deposition contributes to reduced body density.

Given the conserved roles of these genes in neutral lipid hydrolysis, loss of *ces3* and *ces2a* may have shifted the ancestral lipid-metabolism background of Antarctic notothenioids by reducing one route of TAG turnover. This ancestral loss alone does not explain the full range of lipid phenotypes across the clade. Instead, we hypothesize that reduced CES-mediated TAG hydrolysis created a permissive metabolic context that later lineages could exploit to different degrees, particularly in neutrally buoyant or midwater taxa where lipid deposition contributes substantially to reduced body density (**Figure 3D**). This interpretation is consistent with known variation in notothenioid lipid biology, but it remains a testable hypothesis pending direct lipid-turnover measurements across species with contrasting ecologies.

### Ocular water handling and the inactivation of the atypical aquaporin-11 (*aqp11)*

Physiological studies of Antarctic notothenioids have shown that several secreted fluids, including urine, endolymph, and aqueous and vitreous humors, contain little or no antifreeze glycoprotein and have freezing points near −1.0 °C. Because Antarctic seawater is colder, approximately −1.9 °C, these fluids are undercooled *in vivo* and may depend on barriers to ice entry or other mechanisms that prevent nucleation [53]. In vertebrates *AQP11* (Aquaporin-11) gene encodes an atypical intracellular aquaporin expressed in Müller glial cells, where it regulates osmotic water permeability and maintains retinal fluid homeostasis [54–56]. Our comparative genomic analysis revealed inactivating mutations in the *aqp11* gene across all examined cryonotothenioids. In most cryonotothenioids, *aqp11* inactivation was driven by large scale exon deletions, ranging from the complete loss of exon 1 (e.g., *L. nudifrons*, *T. bernacchii*) to the complete deletion of the entire gene in more derived lineages. By contrast, the two *Dissostichus* species retained much of the coding sequence but exhibited multiple frameshift mutations across the first two exons, suggesting lineage-specific sequence decay after ancestral disruption (**Figure 4A**).

**Figure 4.**
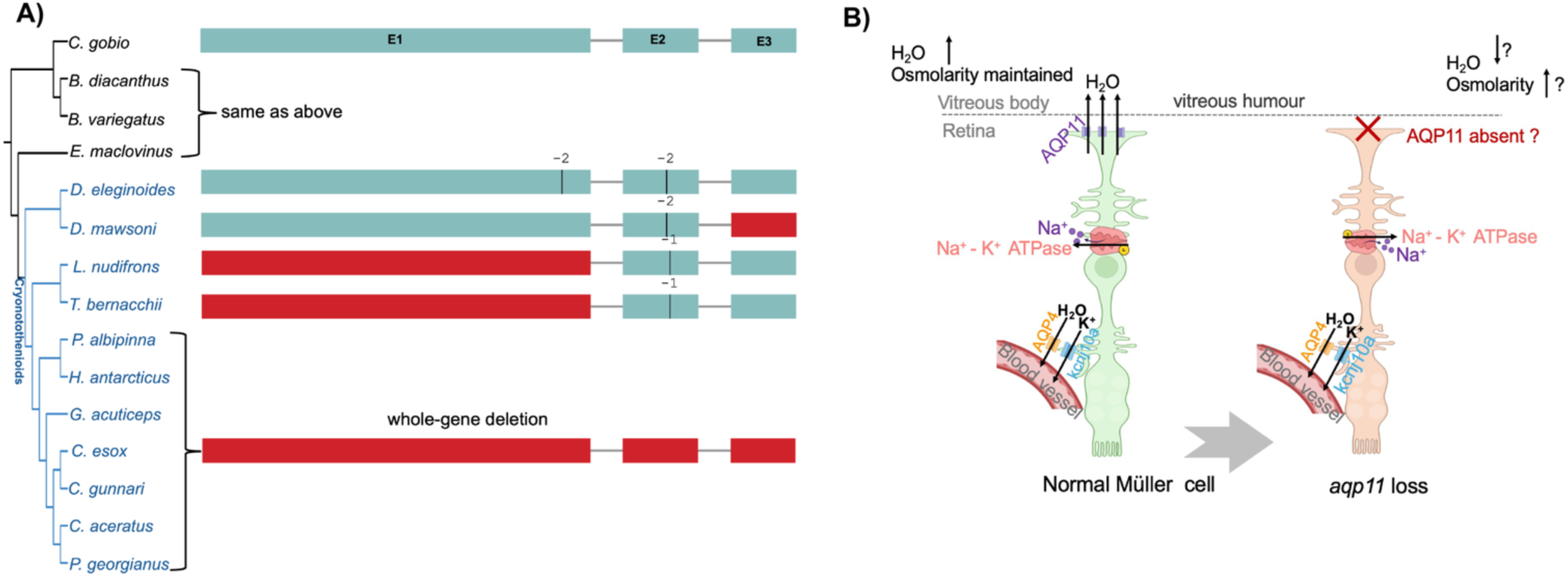
Loss of *aqp11* coding potential and proposed remodeling of ocular/retinal water handling in cryonotothenioids. **A)** *aqp11* gene model comparisons across outgroups and cryonotothenioids. Gene-model colors follow Figure 3. Most cryonotothenioids show exon loss or whole-gene deletion, whereas *Dissostichus* species retain part of the coding sequence but carry frameshifting mutations. **B)** Conceptual model of AQP11-associated water handling at the Müller cell–vitreous interface. In the reference state, AQP11 is shown as part of local Müller glial water transport; after *aqp11* coding loss, local water flux or retinal osmotic balance may be altered. Question marks indicate hypothetical physiological consequences. Arrows indicate water movement, and ion labels indicate associated solute-transport pathways shown for context.

Although the mechanisms underlying maintenance of these undercooled secreted fluids remain unresolved, the loss of *aqp11* raises the possibility that ocular or retinal water handling has been remodeled in cryonotothenioids (**Figure 4B**). This possibility could be tested by comparing AQP11 protein expression and localization in retinal tissues from Antarctic and non-Antarctic notothenioids, together with localized assays of Müller cell water permeability or water transport at the Müller cell–vitreous interface. These experiments would test whether *aqp11* loss is associated with altered retinal water handling, while complementary measurements of ocular fluid osmolality and freezing behavior would be needed to assess consequences for undercooled ocular fluids.

### Loss of *prodhb* and regulation of mitochondrial proline metabolism

In vertebrates, PRODH enzymes catalyze the first step of mitochondrial proline catabolism by oxidizing proline to pyrroline-5-carboxylate (P5C), transferring electrons to ubiquinone and thereby linking proline metabolism to mitochondrial redox state and ATP production [57]. Our comparative genomic analysis identified clade-specific disruption of the *prodhb* (proline dehydrogenase/oxidase 1b) across cryonotothenioids, anchored by a shared deletion of exon 8 and additional lineage-specific frameshifts, whereas this gene remains intact in all four non-Antarctic outgroups (**Figure 5A**).

**Figure 5.**
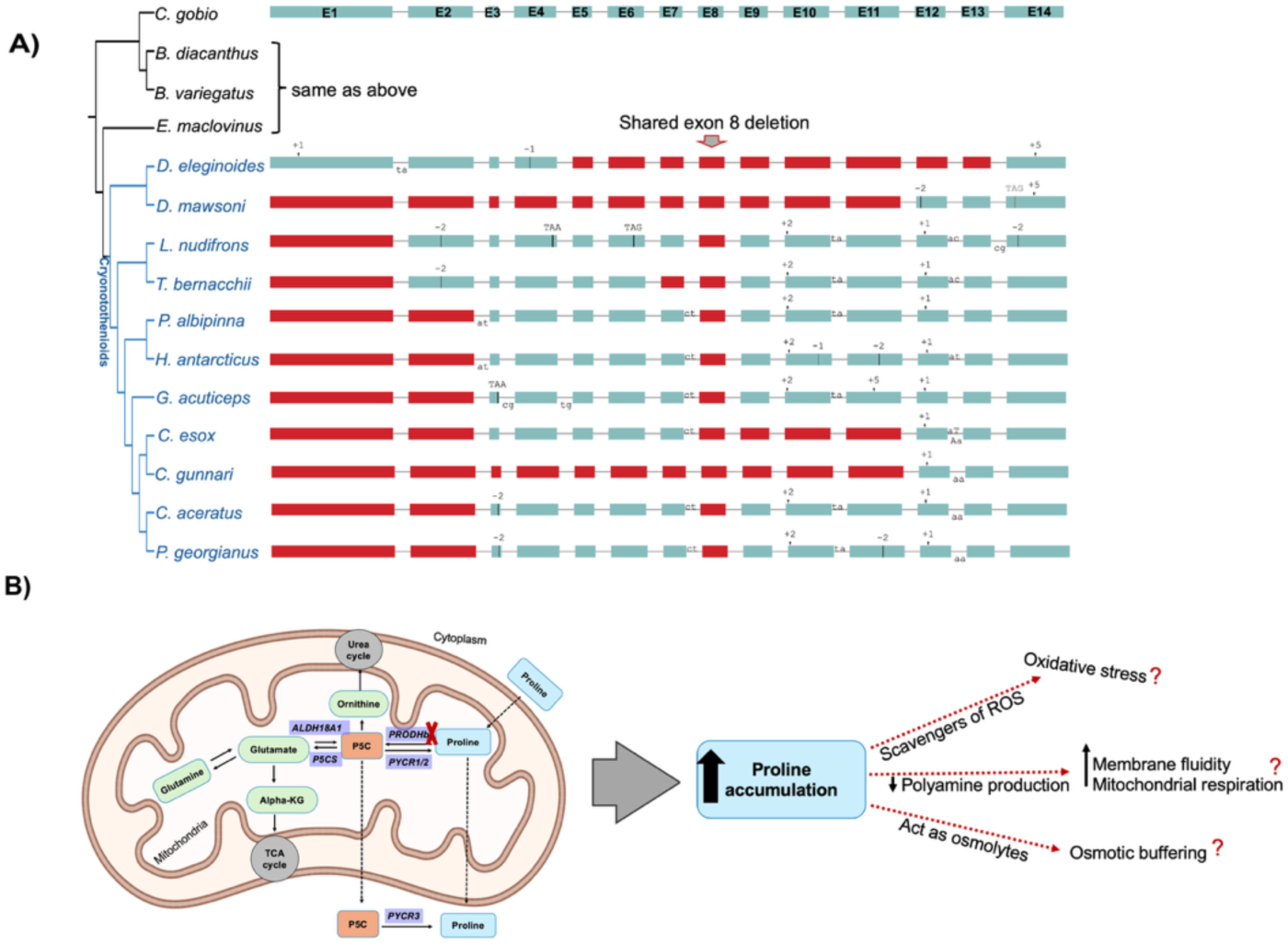
Loss of prodhb and proposed consequences for mitochondrial proline metabolism in cryonotothenioids. **A)** *prodhb* gene model comparisons showing a shared exon 8 deletion in cryonotothenioids, together with additional frameshift and exon-loss events in individual lineages; non-cryonotothenioid outgroups retain intact coding potential. Gene-model colors follow Figure 3. B) Conceptual model of PRODHb in mitochondrial proline oxidation. Loss of *prodhb* is predicted to reduce proline-to-pyrroline-5-carboxylate (P5C) conversion and may alter proline availability, mitochondrial redox balance, and osmotic or cryoprotective stress responses under cold conditions. Question marks indicate proposed effects that require experimental validation.

Because proline can function as a compatible osmolyte and cryoprotectant [58, 59], loss of *prodhb* may alter intracellular proline availability under chronic cold and osmotic stress. Reduced mitochondrial proline oxidation could also affect redox balance by changing electron flow through this pathway [57]. In addition, changes in proline–P5C metabolism may influence downstream amino-acid and stress-response pathways, although links to polyamine metabolism, membrane fluidity, and homeoviscous adaptation remain to be tested directly in notothenioids [60–62]. Together, *prodhb* disruption raises three non-mutually exclusive possibilities: altered osmotic buffering through proline retention, modified redox balance through reduced mitochondrial proline oxidation, and secondary effects on membrane-associated stress responses (**Figure 5B**).

### Loss of *slc5a2* and reduced reliance on glomerular glucose reabsorption

Aglomerular kidneys are widespread among Antarctic notothenioids inhabiting subzero seawater, a renal specialization that reduces or eliminates glomerular filtration and may help limit urinary loss of small peptides and antifreeze glycoproteins [63]. In typical vertebrate kidneys, the *SLC5A2* gene, which encodes sodium-glucose cotransporter 2 (SGLT2), mediates most (∼90%) glucose reabsorption from the glomerular filtrate in the proximal tubule [64, 65]. Our comparative genomic analysis revealed extensive disruption of *slc5a2* across the Antarctic clade, including a large deletion spanning exons 4 through 12, whereas all 15 exons remain intact in non-Antarctic outgroups (**Figure 6A**).

**Figure 6.**
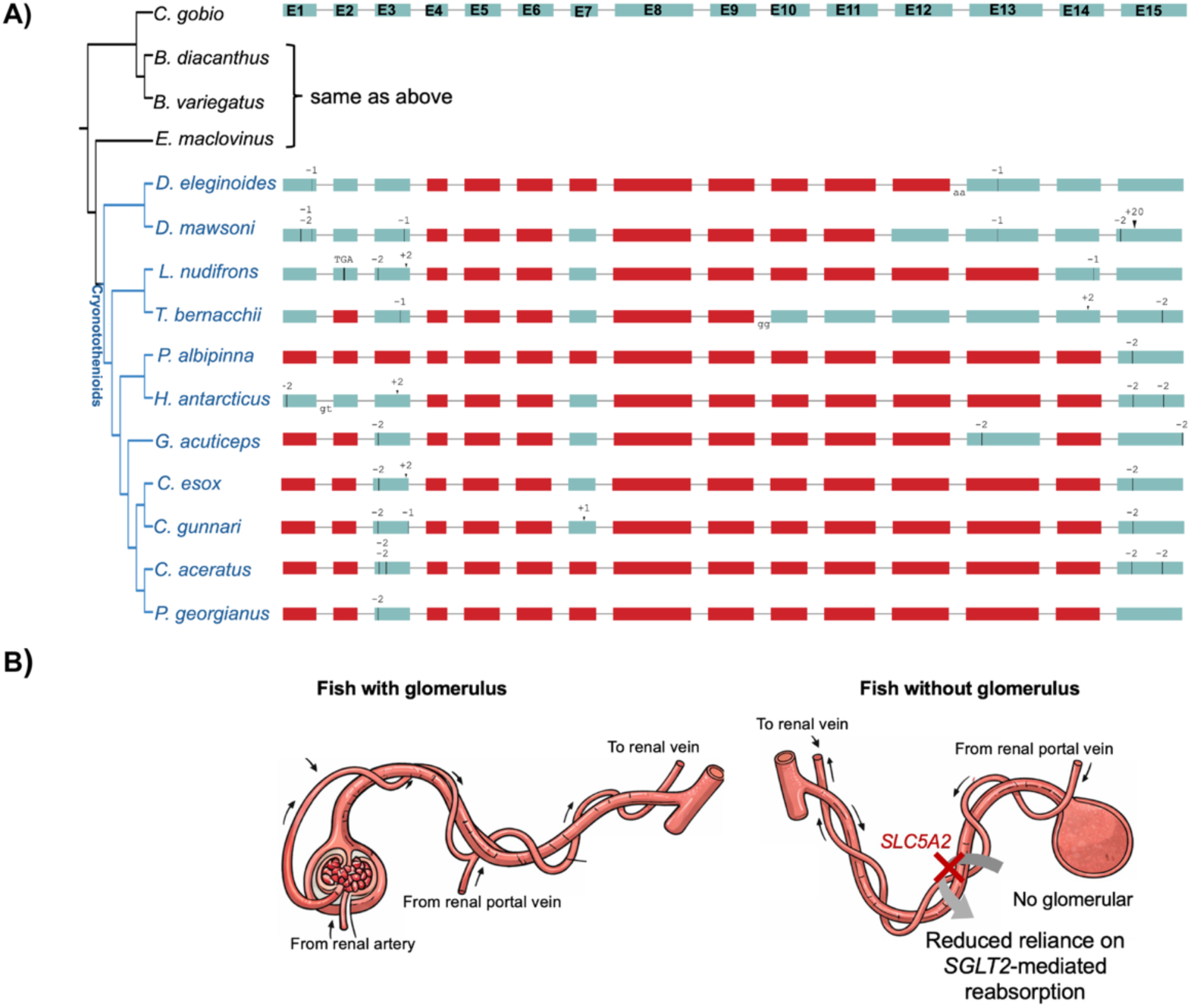
Loss of *slc5a2* and reduced reliance on glomerular glucose reabsorption in cryonotothenioids. **A)** The *slc5a2* gene model comparisons across outgroups and cryonotothenioids. Gene-model colors follow Figure 3. Cryonotothenioids show extensive coding-sequence loss, including a large deletion spanning exons 4–12, whereas non-cryonotothenioid outgroups retain the full exon structure. **B)** Conceptual model linking *slc5a2* loss to reduced dependence on filtrate-based glucose reabsorption. In glomerular kidneys, SLC5A2/SGLT2 mediates glucose reabsorption from the glomerular filtrate in the proximal tubule. In fishes without glomerular filtration, the filtered glucose load is reduced, which may relax selective pressure on *slc5a2*-mediated glucose recovery.

Because *slc5a2* acts on glucose recovered from glomerular filtrate, its loss may reflect reduced selective pressure to maintain proximal-tubule glucose reabsorption in lineages with reduced or absent filtration. We also identified loss of coding potential in *abrab* (actin binding Rho activating protein b), also known as *abra*, a candidate renal-development-associated gene implicated in zebrafish pronephros development [66]. Collectively, these results suggest that losses affecting renal filtration or glucose reabsorption pathways may have accompanied renal specialization in Antarctic notothenioids (**Figure 6B)**.

### Loss of *bglap* and skeletal mineralization

Antarctic notothenioids exhibit reduced skeletal mass and mineralization relative to many temperate fishes, a trait associated with reduced body density and buoyancy in lineages that lack a swim bladder [67–70]. In vertebrates, *BGLAP* encodes osteocalcin, a late marker of osteoblast differentiation and an important component of bone formation [71]. Knockdown of *BGLAP* in human cell lines reduces overall mineralization, and *bglap* expression is downregulated in zebrafish models of skeletal disorders [72, 73]. Consistent with this low-density skeletal phenotype, our comparative genomic analysis revealed the degradation of *bglap* across the examined Antarctic species. In seven of eleven Antarctic species, *bglap* is completely deleted from the genome assemblies, whereas in *Dissostichus mawsoni* we identified a 17-bp frameshifting insertion predicted to cause premature termination (**Figure 7A**).

**Figure 7.**
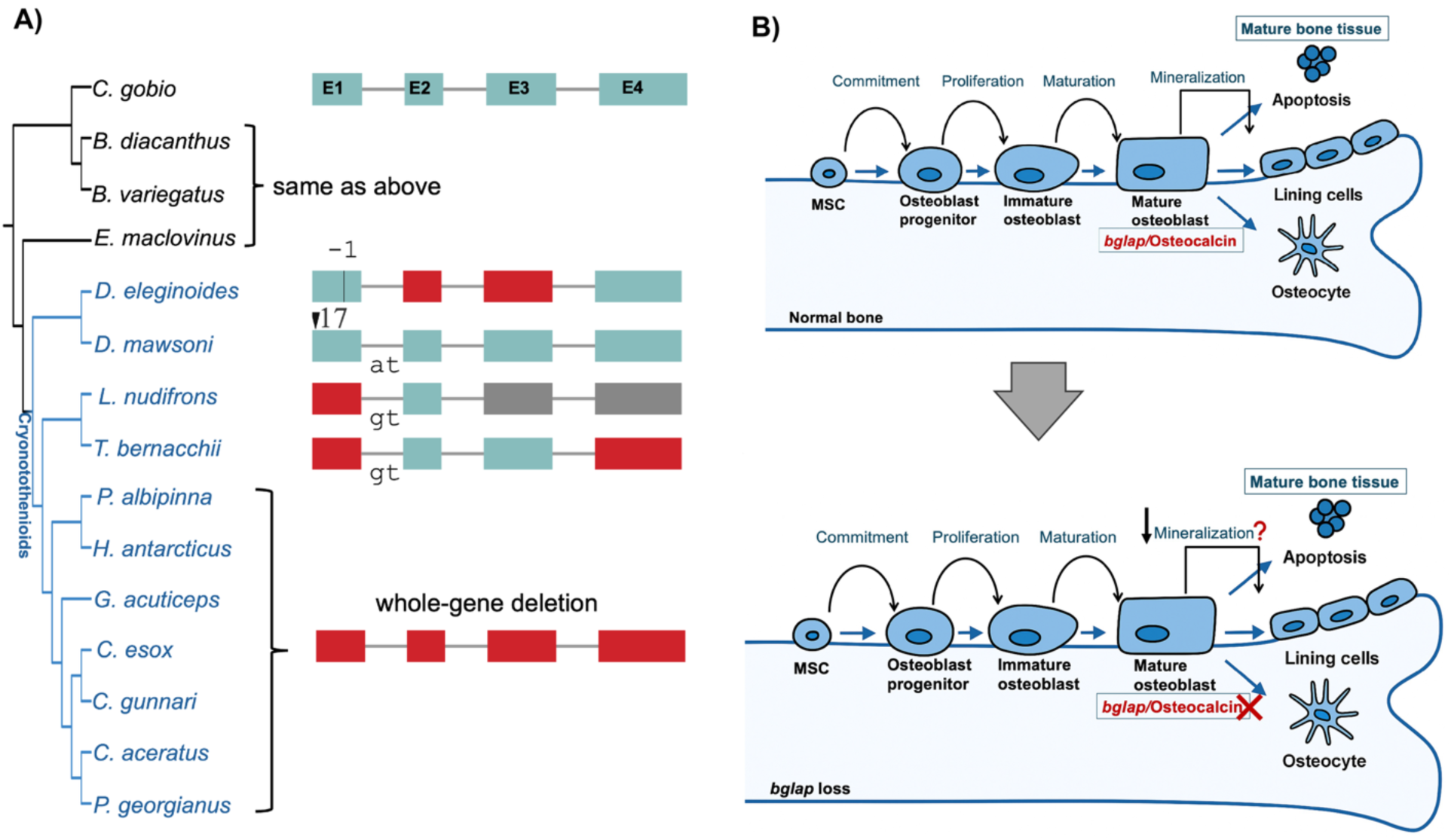
Loss of *bglap* coding potential and a proposed link to skeletal mineralization in cryonotothenioids. A). *bglap* gene model comparisons across non-cryonotothenioid outgroups and cryonotothenioids. Gene-model colors follow Figure 3. Non-cryonotothenioid outgroups retain an intact four-exon structure, whereas several cryonotothenioids show complete or near-complete loss of *bglap* coding sequence. In Dissostichus mawsoni, a 17-bp frameshifting insertion is predicted to disrupt the coding sequence. **B).** Conceptual model showing the role of BGLAP/osteocalcin during late osteoblast maturation and bone mineralization. Loss of *bglap* coding potential may alter osteoblast maturation or reduce skeletal mineralization in cryonotothenioids.

Reduced skeletal density in cryonotothenioids has been linked to heterochronic and pedomorphic skeletal features, including reduced ossification and retention of cartilage or juvenile skeletal features into adulthood [74]. In this context, loss of *bglap* may have contributed to altered osteoblast maturation or reduced skeletal mineralization in Antarctic notothenioids (**Figure 7B**). Notably, the paralog *bglapl* (bone gamma-carboxyglutamic acid protein-like or osteocalcin-2) remains intact across the Antarctic clade, raising the possibility that osteocalcin-related functions are partially retained or compensated. The extent to which *bglapl* buffers the loss of *bglap* during bone remodeling remains an open question.

### Coordinated disruption of multiple tRNA modification genes involved in wybutosine biosynthesis pathway

In most eukaryotes, the wybutosine (yW) biosynthesis pathway modifies phenylalanine tRNA (tRNA^Phe^) at position 37, stabilizing codon–anticodon interactions and helping maintain reading-frame fidelity during translation [75]. This multi-step pathway involves TRM5 and the TYW complex (TYW1-TYW5) (**Figure 8B**). Our comparative genomic analysis revealed the clade-wide pseudogenization of four sequential genes in this cascade—*tyw2*, *tyw3*, *tyw4*, and *tyw5*—across all examined cryonotothenioids. While all these loci remain fully intact across all four non-Antarctic outgroups, the Antarctic clade exhibits extensive coding-sequence decay with a shared deletion of exons 4 through 6 in *tyw2* (**Figure 8A)**, and additional frameshifts or exon deletions in downstream pathway genes **(Figure 8B, Figures S1–S3)**.

**Figure 8.**
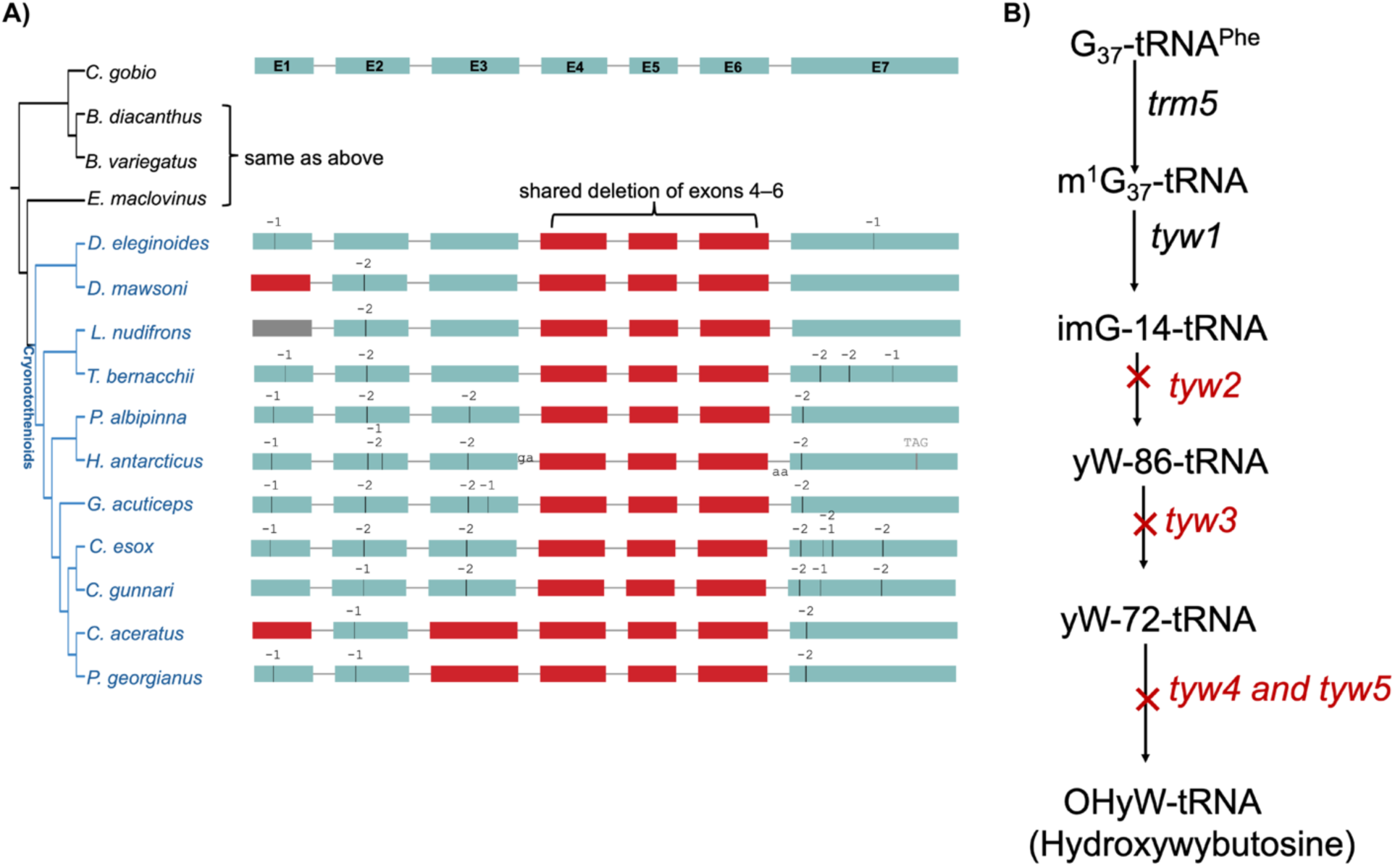
Coordinated loss of wybutosine biosynthesis genes in cryonotothenioids. **A)** *tyw2* gene model comparisons across non-cryonotothenioid outgroups and cryonotothenioids. Gene-model colors follow Figure 3. Cryonotothenioids show a shared deletion spanning multiple internal exons of *tyw2*, together with additional lineage-specific coding-sequence changes, whereas outgroups retain the intact exon structure. **B)** Simplified wybutosine (yW) biosynthesis pathway showing sequential modification of G37-tRNA^Phe^ to hydroxywybutosine. Genes shown in red with red crosses indicate yW-pathway genes inferred to have lost coding potential in cryonotothenioids, including *tyw2, tyw3, tyw4, and tyw5*. Black gene labels indicate upstream pathway components not inferred as lost in this analysis.

The coordinated disruption of multiple sequential enzymes suggests that canonical yW biosynthesis has been strongly reduced or lost in the Antarctic clade. Experimental work in eukaryotic systems has shown that yW modification and its biosynthetic intermediates influence ribosomal frameshifting [75], raising the possibility that loss/reduce of yW biosynthesis could alter translational fidelity in cryonotothenioids. Antarctic fishes show unusual features of protein metabolism, including differences in protein synthesis, protein retention, and RNA-to-protein ratios relative to temperate fishes [76]. Whether these broader protein-metabolism traits interact with yW-pathway loss remains unknown; direct assays of tRNA modification status, ribosomal frameshifting, and translation fidelity under cold conditions will be required to test the functional consequences of this pathway-level loss.

### Loss of core clock genes and remodeling of circadian regulation

Circadian rhythms coordinate daily oscillations in physiology and behavior through endogenous molecular clocks entrained by environmental cues [77]. In Antarctic notothenioids, we identified complete exon deletion of five circadian-related genes. Four of these genes belong to the core *cryptochrome* and *period* families (*cry1b*, *cry2*, *per2a*, and *per3*). We also identified loss of *fbxl3l* (F-box and leucine-rich repeat protein 3-like), a circadian-associated F-box gene whose mammalian homolog regulates clock timing by promoting CRY protein ubiquitination and turnover [78], representing an additional circadian-regulatory loss in this study. Previous work reported apparent *per2a* and *per3* pseudogenization in the non-Antarctic outgroup *C. gobio* [23, 24], and our analysis also recovered a multi-exon deletion disrupting *per2a* and *per3* in this species. Because our dataset reveals that both *per2a* and *per3* remain intact in other two *Bovichtidae* species, the *C. gobio* loss and the clade-wide loss observed across Antarctic notothenioids are best interpreted as independent events.

Recent comparative genomic studies have reported erosion of circadian gene repertoires in polar and high-latitude fishes, particularly among *cry* and *per* paralogs [23, 24, 79]. The losses identified here extend this pattern across Antarctic notothenioids and suggest remodeling of canonical clock pathways under extreme seasonal photoperiods. Because at least one *cry* and one *per* paralog remain intact across the clade [24], these data do not imply complete loss of circadian regulation. Instead, they point to a modified circadian system whose functional consequences remain unresolved. Future behavioral, expression, and physiological assays under polar light regimes will be needed to determine whether remaining rhythms are primarily light-entrained or are also influenced by non-photic cues such as tidal cycles or seasonal feeding patterns.

### Loss of neuropeptide gene *cart2* and appetite-regulatory networks

In teleost fishes, appetite regulation and energy homeostasis are controlled by interacting central neuroendocrine pathways. Within this network, *cart2* (cocaine- and amphetamine-regulated transcript 2) encodes a conserved anorexigenic neuropeptide involved in feeding suppression and energy-balance regulation [80, 81]. Our comparative genomic analysis revealed truncation of *cart2* across Antarctic notothenioids. In most Antarctic species, *cart2* loss involved deletion of all three exons, whereas in *Dissostichus* one exon remains detectable (**Figure S4**).

Given the anorexigenic role of CART peptides, loss of *cart2* may remove one satiety-related signal and reshape appetite-regulatory networks in Antarctic notothenioids. The net physiological effect is likely to depend on compensation by other *cart* paralogs and interactions with additional anorexigenic and orexigenic pathways, including *pomc*, *npy*, and orexin-related signaling [81]. Targeted feeding assays and expression analyses will be needed to determine how *cart2* loss affects appetite regulation in this clade.

### Recovery of known hemoglobin gene loss and identification of *vps4a* coding-sequence loss in Antarctic icefishes

A hallmark of Antarctic icefishes is the loss of mature circulating erythrocytes and functional hemoglobin, a phenotype associated with well-characterized erosion of globin loci in Channichthyidae [25–28, 82, 83]. Our pipeline recovered this expected icefish-specific signal: all examined icefishes lacked several hemoglobin genes, including five α-globin genes (*hba1, hba2, hbae1.2, hbae4,* and *hbae5*) and two β-globin genes (*hbbe2* and *hbbe3*), whereas these loci were retained in red-blooded notothenioids and non-Antarctic outgroups. Recovery of this well-established gene-loss case supports the ability of our pipeline to detect known lineage-specific losses and provides context for interpreting additional, less-characterized icefish-specific candidates.

Beyond globin loci, we identified loss of coding potential in *vps4a* across all examined icefishes. Red-blooded notothenioids and all four non-Antarctic outgroups retain an intact *vps4a* coding sequence, whereas icefishes show extensive coding-region disruption, including multi-exon deletions and frameshift mutations predicted to abolish the open reading frame (**Figure 9A**). *Vps4a* encodes an AAA-ATPase involved in disassembly of the ESCRT (Endosomal Sorting Complex Required for Transport) machinery, a process essential for membrane remodeling, vesicular sorting, cytokinesis, and erythropoiesis [84].

**Figure 9.**
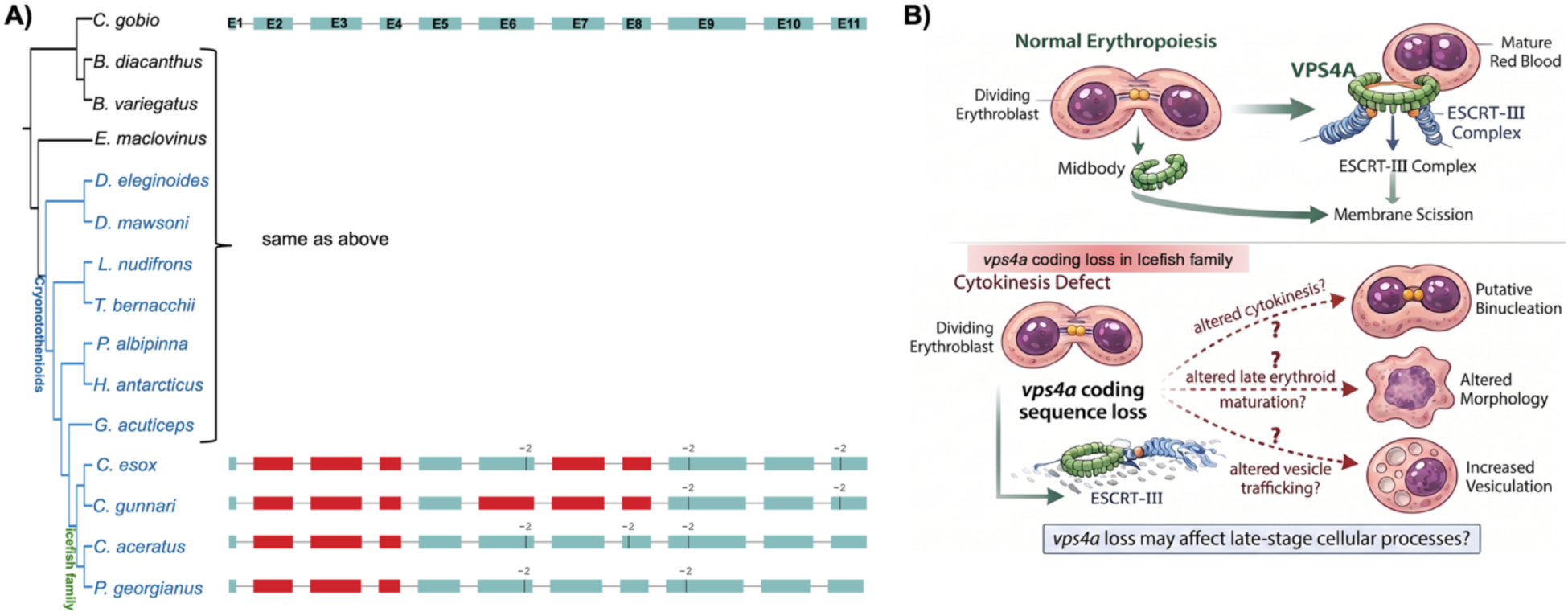
Icefish-specific loss of *vps4a* coding potential and proposed effects on ESCRT-dependent cellular processes. **A)** *vps4a* gene model comparisons across outgroups, red-blooded cryonotothenioids, and icefishes. Gene-model colors follow Figure 3. Red-blooded cryonotothenioids and non-cryonotothenioid outgroups retain intact *vps4a* coding sequence, whereas icefishes show extensive coding-region disruption, including exon deletions and frameshifting mutations. **B)** Conceptual model of VPS4A function in ESCRT-mediated membrane scission during cell division. In the reference state, VPS4A promotes ESCRT-III disassembly during cytokinesis and membrane remodeling. In icefishes, *vps4a* coding-sequence loss may alter ESCRT-dependent processes, including cytokinesis, vesicle trafficking, and late erythroid cell maturation.

Recent studies demonstrate that despite lacking circulating erythrocytes, icefish retain early erythroid progenitors in hematopoietic tissue, indicating that erythropoiesis fails at a later developmental stage [82]. In this context, loss of *vps4a* coding potential raises the possibility that ESCRT-dependent membrane remodeling or cytokinesis pathways were altered during icefish erythroid evolution. This interpretation is compatible with reduced constraint on erythroid-associated functions after loss of hemoglobin-based oxygen transport and mature erythrocytes (**Figure 9B**), but alternative explanations, including compensation by other ESCRT components in non-erythroid tissues, remain possible. Comparative analyses of ESCRT-related trafficking, cytokinesis, and erythroid cell development will be needed to clarify the functional consequences of *vps4a* loss in icefishes.

## Discussion

We systematically identified gene losses shared across the Antarctic notothenioid clade and additional losses restricted to the derived icefish lineage. Many of the shared gene losses most parsimoniously trace to the common ancestor of the Cryonotothenioidea and affect diverse biological systems, including lipid and amino acid metabolism, water transport, renal glucose reabsorption, bone mineralization, wybutosine/tRNA modification, circadian regulation, and appetite control. These losses provide candidate genomic links to several well-known Antarctic notothenioid traits, including lineage-variable lipid storage, freezing avoidance, reduced or absent glomerular filtration, reduced skeletal mineralization, and altered circadian regulation. In icefishes, we recovered the well-established loss of hemoglobin genes and identified additional lineage-specific losses, including *vps4a* and other hematological or vesicular-trafficking candidates. Together, these results suggest that gene loss is not an isolated feature of Antarctic notothenioid evolution, but a recurrent component of genome remodeling under chronic cold.

Many genes lost in Antarctic notothenioids are associated with disease phenotypes in humans and model vertebrates, making them a compelling system for studying naturally occurring gene loss. In laboratory systems, engineered loss-of-function mutations often produce severe or early developmental phenotypes. In contrast, the losses described here have persisted in viable wild populations over evolutionary time. This contrast provides an opportunity to ask how ecological setting and genomic background alter the consequences of gene loss. In this sense, Antarctic notothenioids can be viewed as evolutionary mutant models: natural systems in which gene losses resembling engineered knockouts are embedded within ecological and genomic contexts that permit organismal viability.

Representative examples span metabolic, renal, skeletal, circadian, and hematological systems. Loss of *prodhb* intersects with proline-catabolism disorders such as hyperprolinemia [85], *slc5a2* loss relates to renal glucose transport and familial renal glucosuria [86, 87], and *bglap* loss connects to skeletal mineralization phenotypes summarized in Table 1 [73]. Likewise, losses of *per/cry* clock components intersect with circadian disorders, including sleep-phase disorders and seasonal mood phenotypes [88, 89], whereas icefish-specific *vps4a* loss points to ESCRT-dependent pathways associated with congenital dyserythropoietic anemia, neurodevelopmental disease, and zebrafish sensorimotor defects [90, 91]. These examples extend the significance of Antarctic notothenioid gene loss beyond cold adaptation alone by highlighting naturally fixed coding-gene losses in pathways that are disease-associated in other vertebrates.

A central question raised by these results is why losses that are deleterious or pleiotropic in other vertebrates are apparently tolerated in Antarctic notothenioids. Many of the affected genes act in multiple tissues or developmental contexts, so their disruption in humans or model organisms can produce broad downstream effects rather than a single isolated phenotype. In Antarctic notothenioids, comparable negative effects may be absent, environmentally buffered, compensated by paralogs, or present only as subtle traits that have not yet been measured. For example, the cold, oxygen-rich Southern Ocean buffers the loss of hemoglobin-based oxygen transport in icefishes [25, 92], while intact paralogs or alternative pathways may buffer losses affecting bone mineralization, appetite regulation, circadian timing, translation, or vesicular trafficking. These patterns motivate closer physiological examination of notothenioids, not only to identify hidden consequences of gene loss, but also to understand how natural systems mitigate the negative pleiotropic effects that similar losses produce in other vertebrate contexts.

More broadly, these findings support a systems-level view of gene loss in Antarctic notothenioids. Individual losses may have arisen through relaxed constraint, neutral decay, compensation, or adaptive change, or combinations of these processes. Once fixed, shared losses become part of the inherited genomic background on which later evolution occurs. In Antarctic notothenioids, long-term thermal stability, high oxygen availability, and the absence or reduction of selective pressures present in other environments may have permitted the persistence of losses that would be costly in other vertebrate contexts. This genome-wide catalog therefore provides a framework for testing how gene loss reshapes physiological systems under constant cold.

## Conclusion

Our analyses move gene loss in Antarctic notothenioids beyond isolated examples and into a testable comparative framework. By combining conservative gene-loss calls with phylogenetic context, we nominate a set of naturally occurring coding-gene losses whose persistence in viable wild populations raises important questions about environmental buffering, compensation, and pathway rewiring. The functional links proposed here are based largely on orthology and evidence from model vertebrates; their physiological consequences in notothenioids remain to be tested directly. Future work should combine comparative expression analyses, targeted physiological measurements, and functional assays to determine which losses are neutral, tolerated, compensatory, or adaptive. Such studies will clarify how gene loss reshapes vertebrate physiology under chronic cold and may reveal natural mechanisms that buffer disease-associated pathways in wild populations.

## Declarations

### Ethics approval and consent to participate

Not applicable.

## Consent for publication

Not applicable.

## Availability of data and materials

The publicly available genome assemblies analyzed during the current study are available from NCBI GenBank/RefSeq under the accession numbers listed in Additional file 1: Table S1. The publicly available RNA-seq datasets analyzed during the current study are available from the NCBI Sequence Read Archive under the BioProject or SRR accession numbers listed in Additional file 2: Table S2. The datasets generated during the current study will be deposited in Figshare prior to publication.

## Authors’ contributions

VL and XZ conceived and designed the study. VL curated the data, performed the analyses, generated the figures and tables, and drafted the manuscript. AJA and JMD contributed to data interpretation and critically revised the manuscript. XZ supervised the project, acquired funding, provided resources, contributed to interpretation, and critically revised the manuscript. All authors read and approved the final manuscript.

## Acknowledgements

We thank Professor Yizhi Sun, Christian Tipsmark, and Isabelle R. Miousse for valuable feedback. We also acknowledge the Arkansas High Performance Computing Center which is funded through multiple National Science Foundation grants and the Arkansas Economic Development Commission.

## References

1. Blomme T, Vandepoele K, De Bodt S, Simillion C, Maere S, Van de Peer Y: The gain and loss of genes during 600 million years of vertebrate evolution. Genome biology 2006, 7(5):R43.

2. Fernández R, Gabaldón T: Gene gain and loss across the metazoan tree of life. Nat Ecol Evol 4: 524–533. In.; 2020.

3. Albalat R, Cañestro C: Evolution by gene loss. Nature Reviews Genetics 2016, 17(7):379–391.

4. Hirsh AE, Fraser HB: Protein dispensability and rate of evolution. Nature 2001, 411(6841):1046–1049.

5. Pál C, Papp B, Lercher MJ: An integrated view of protein evolution. Nature reviews genetics 2006, 7(5):337–348.

6. Sharma V, Hecker N, Roscito JG, Foerster L, Langer BE, Hiller M: A genomics approach reveals insights into the importance of gene losses for mammalian adaptations. Nature communications 2018, 9(1):1215.

7. Olson MV: When less is more: gene loss as an engine of evolutionary change. The American Journal of Human Genetics 1999, 64(1):18–23.

8. Helsen J, Voordeckers K, Vanderwaeren L, Santermans T, Tsontaki M, Verstrepen KJ, Jelier R: Gene loss predictably drives evolutionary adaptation. Molecular Biology and Evolution 2020, 37(10):2989–3002.

9. Miyata T, Yasunaga T, Nishida T: Nucleotide sequence divergence and functional constraint in mRNA evolution. Proceedings of the National Academy of Sciences 1980, 77(12):7328–7332.

10. Krylov DM, Wolf YI, Rogozin IB, Koonin EV: Gene loss, protein sequence divergence, gene dispensability, expression level, and interactivity are correlated in eukaryotic evolution. Genome research 2003, 13(10):2229.

11. Jordan IK, Rogozin IB, Wolf YI, Koonin EV: Essential genes are more evolutionarily conserved than are nonessential genes in bacteria. Genome research 2002, 12(6):962–968.

12. Yang J, Gu Z, Li W-H: Rate of Protein Evolution vs. Fitness Effect of Gene Deletion. Mol Biol Evol 20 (5), 772–774 2003.

13. Bista I, Wood JM, Desvignes T, McCarthy SA, Matschiner M, Ning Z, Tracey A, Torrance J, Sims Y, Chow W: Genomics of cold adaptations in the Antarctic notothenioid fish radiation. Nature Communications 2023, 14(1):3412.

14. Near TJ, Dornburg A, Kuhn KL, Eastman JT, Pennington JN, Patarnello T, Zane L, Fernández DA, Jones CD: Ancient climate change, antifreeze, and the evolutionary diversification of Antarctic fishes. Proceedings of the National Academy of Sciences 2012, 109(9):3434–3439.

15. Beers JM, Jayasundara N: Antarctic notothenioid fish: what are the future consequences of ‘losses’ and ‘gains’ acquired during long-term evolution at cold and stable temperatures? The Journal of Experimental Biology 2015, 218(12):1834–1845.

16. Eastman JT: The nature of the diversity of Antarctic fishes. Polar biology 2005, 28(2):93–107.

17. Auvinet J, Graça P, Dettai A, Amores A, Postlethwait JH, Detrich III HW, Ozouf-Costaz C, Coriton O, Higuet D: Multiple independent chromosomal fusions accompanied the radiation of the Antarctic teleost genus Trematomus (Notothenioidei: Nototheniidae). BMC Evolutionary Biology 2020, 20(1):39.

18. Amores A, Wilson CA, Allard CA, Detrich III HW, Postlethwait JH: Cold fusion: massive karyotype evolution in the Antarctic bullhead notothen Notothenia coriiceps. G3: Genes, Genomes, Genetics 2017, 7(7):2195–2207.

19. Devries AL: Glycoproteins as biological antifreeze agents in Antarctic fishes. Science 1971, 172(3988):1152–1155.

20. Hofmann GE, Buckley BA, Airaksinen S, Keen JE, Somero GN: Heat-shock protein expression is absent in the Antarctic fish Trematomus bernacchii (family Nototheniidae). Journal of Experimental Biology 2000, 203(15):2331–2339.

21. Buckley BA, Place SP, Hofmann GE: Regulation of heat shock genes in isolated hepatocytes from an Antarctic fish, Trematomus bernacchii. Journal of Experimental Biology 2004, 207(21):3649–3656.

22. Beers JM, Sidell BD: Thermal tolerance of Antarctic notothenioid fishes correlates with level of circulating hemoglobin. Physiological and Biochemical Zoology 2011, 84(4):353–362.

23. Cheng C-HC, Rivera-Colón AG, Minhas BF, Wilson L, Rayamajhi N, Vargas-Chacoff L, Catchen JM: Chromosome-level genome assembly and circadian gene repertoire of the patagonia Blennie Eleginops maclovinus—the closest ancestral proxy of Antarctic Cryonotothenioids. Genes 2023, 14(6):1196.

24. Wright DB, Zhang Y, Daane JM: Convergent latitudinal erosion of circadian systems in a rapidly diversifying order of fishes. bioRxiv 2025.

25. Ruud JT: Vertebrates without erythrocytes and blood pigment. Nature 1954, 173(4410):848–850.

26. di Prisco G, Cocca E, Parker SK, Detrich III HW: Tracking the evolutionary loss of hemoglobin expression by the white-blooded Antarctic icefishes. Gene 2002, 295(2):185–191.

27. Near TJ, Parker SK, Detrich III HW: A genomic fossil reveals key steps in hemoglobin loss by the Antarctic icefishes. Molecular Biology and Evolution 2006, 23(11):2008–2016.

28. Sidell BD, O’Brien KM: When bad things happen to good fish: the loss of hemoglobin and myoglobin expression in Antarctic icefishes. Journal of Experimental Biology 2006, 209(10):1791–1802.

29. Daane JM, Dornburg A, Smits P, MacGuigan DJ, Brent Hawkins M, Near TJ, William Detrich III H, Harris MP: Historical contingency shapes adaptive radiation in Antarctic fishes. Nature Ecology & Evolution 2019, 3(7):1102–1109.

30. Shin SC, Kim S, Kim H-W, Lee JH, Kim J-H: Gene loss in Antarctic icefish: evolutionary adaptations mimicking Fanconi Anemia? BMC genomics 2024, 25(1):1102.

31. Balushkin A: Morphology, classification, and evolution of notothenioid fishes of the Southern Ocean (Notothenioidei, Perciformes). Journal of Ichthyology 2000(suppl. 1).

32. Eastman JT, Eakin RR: Checklist of the species of notothenioid fishes. Antarctic Science 2021, 33(3):273–280.

33. Simão FA, Waterhouse RM, Ioannidis P, Kriventseva EV, Zdobnov EM: BUSCO: assessing genome assembly and annotation completeness with single-copy orthologs. Bioinformatics 2015, 31(19):3210–3212.

34. Bolger AM, Lohse M, Usadel B: Trimmomatic: a flexible trimmer for Illumina sequence data. Bioinformatics 2014, 30(15):2114–2120.

35. Grabherr MG, Haas BJ, Yassour M, Levin JZ, Thompson DA, Amit I, Adiconis X, Fan L, Raychowdhury R, Zeng Q: Full-length transcriptome assembly from RNA-Seq data without a reference genome. Nature biotechnology 2011, 29(7):644–652.

36. Dobin A, Davis CA, Schlesinger F, Drenkow J, Zaleski C, Jha S, Batut P, Chaisson M, Gingeras TR: STAR: ultrafast universal RNA-seq aligner. Bioinformatics 2013, 29(1):15–21.

37. Emms DM, Kelly S: OrthoFinder: phylogenetic orthology inference for comparative genomics. Genome biology 2019, 20(1):238.

38. Edgar RC: MUSCLE: a multiple sequence alignment method with reduced time and space complexity. BMC bioinformatics 2004, 5(1):113.

39. Nguyen L-T, Schmidt HA, Von Haeseler A, Minh BQ: IQ-TREE: a fast and effective stochastic algorithm for estimating maximum-likelihood phylogenies. Molecular biology and evolution 2015, 32(1):268–274.

40. Emms D, Kelly S: STAG: species tree inference from all genes. BioRxiv 2018:267914.

41. Near TJ: Estimating divergence times of notothenioid fishes using a fossil-calibrated molecular clock. Antarctic Science 2004, 16(1):37–44.

42. Suarez HG, Langer BE, Ladde P, Hiller M: chainCleaner improves genome alignment specificity and sensitivity. Bioinformatics 2017, 33(11):1596–1603.

43. Osipova E, Hecker N, Hiller M: RepeatFiller newly identifies megabases of aligning repetitive sequences and improves annotations of conserved non-exonic elements. Gigascience 2019, 8(11):giz132.

44. Kirilenko BM, Munegowda C, Osipova E, Jebb D, Sharma V, Blumer M, Morales AE, Ahmed A-W, Kontopoulos D-G, Hilgers L: Integrating gene annotation with orthology inference at scale. Science 2023, 380(6643):eabn3107.

45. Camacho C, Coulouris G, Avagyan V, Ma N, Papadopoulos J, Bealer K, Madden TL: BLAST+: architecture and applications. BMC bioinformatics 2009, 10(1):421.

46. Reimand J, Kull M, Peterson H, Hansen J, Vilo J: g: Profiler—a web-based toolset for functional profiling of gene lists from large-scale experiments. Nucleic acids research 2007, 35(suppl_2):W193–W200.

47. Eastman JT, DeVries AL: Buoyancy studies of notothenioid fishes in McMurdo Sound, Antarctica. Copeia 1982:385–393.

48. Clarke A, Doherty N, DeVries A, Eastman J: Lipid content and composition of three species of Antarctic fish in relation to buoyancy. Polar Biology 1984, 3(2):77–83.

49. Phleger C, Nichols PD, Erb E, Williams R: Lipids of the notothenioid fishes Trematomus spp. and Pagothenia borchgrevinki from East Antarctica. Polar Biology 1999, 22(4):241–247.

50. Wei E, Ali YB, Lyon J, Wang H, Nelson R, Dolinsky VW, Dyck JR, Mitchell G, Korbutt GS, Lehner R: Loss of TGH/Ces3 in mice decreases blood lipids, improves glucose tolerance, and increases energy expenditure. Cell metabolism 2010, 11(3):183–193.

51. Li Y, Zalzala M, Jadhav K, Xu Y, Kasumov T, Yin L, Zhang Y: Carboxylesterase 2 prevents liver steatosis by modulating lipolysis, endoplasmic reticulum stress, and lipogenesis and is regulated by hepatocyte nuclear factor 4 alpha in mice. Hepatology 2016, 63(6):1860–1874.

52. Lian J, Nelson R, Lehner R: Carboxylesterases in lipid metabolism: from mouse to human. Protein & cell 2018, 9(2):178–195.

53. Ahlgren JA, Cheng C-HC, Schrag JD, Devries AL: Freezing avoidance and the distribution of antifreeze glycopeptides in body fluids and tissues of Antarctic fish. Journal of Experimental Biology 1988, 137(1):549–563.

54. Ikeda M, Andoo A, Shimono M, Takamatsu N, Taki A, Muta K, Matsushita W, Uechi T, Matsuzaki T, Kenmochi N: The NPC motif of aquaporin-11, unlike the NPA motif of known aquaporins, is essential for full expression of molecular function. Journal of Biological Chemistry 2011, 286(5):3342–3350.

55. Deeg CA, Amann B, Lutz K, Hirmer S, Lutterberg K, Kremmer E, Hauck SM: Aquaporin 11, a regulator of water efflux at retinal Müller glial cell surface decreases concomitant with immune-mediated gliosis. Journal of neuroinflammation 2016, 13(1):89.

56. Zhang C, Luo D, Xie H, Yang Q, Liu D, Tang L, Zhang J, Li W, Tian H, Lu L: Aquaporin 11 alleviates retinal Müller intracellular edema through water efflux in diabetic retinopathy. Pharmacological Research 2023, 187:106559.

57. Pallag G, Nazarian S, Ravasz D, Bui D, Komlódi T, Doerrier C, Gnaiger E, Seyfried TN, Chinopoulos C: Proline oxidation supports mitochondrial ATP production when complex I is inhibited. International journal of molecular sciences 2022, 23(9):5111.

58. Dou M, Lu C, Sun Z, Rao W: Natural cryoprotectants combinations of l-proline and trehalose for red blood cells cryopreservation. Cryobiology 2019, 91:23–29.

59. Liu X, Pan Y, Liu F, He Y, Zhu Q, Liu Z, Zhan X, Tan S: A review of the material characteristics, antifreeze mechanisms, and applications of cryoprotectants (CPAs). Journal of nanomaterials 2021, 2021(1):9990709.

60. Roberts D, Dumbroff E, Thompson J: Exogenous polyamines alter membrane fluidity in bean leaves—a basis for potential misinterpretation of their true physiological role. Planta 1986, 167(3):395–401.

61. Zheliaskova A, Naydenova S, Petrov A: Interaction of phospholipid bilayers with polyamines of different length. European Biophysics Journal 2000, 29(2):153–157.

62. Hazel JR: Thermal adaptation in biological membranes: is homeoviscous adaptation the explanation? Annual review of physiology 1995, 57(1):19–42.

63. Dobbs III GH, Lin Y, DeVries AL: Aglomerularism in Antarctic fish. Science 1974, 185(4153):793–794.

64. Wright EM, Turk E: The sodium/glucose cotransport family SLC5. Pflügers Archiv 2004, 447(5):510–518.

65. Wright EM: Renal Na+-glucose cotransporters. American Journal of Physiology-Renal Physiology 2001, 280(1):F10–F18.

66. Burghardt T, Kastner J, Suleiman H, Rivera-Milla E, Stepanova N, Lottaz C, Kubitza M, Böger CA, Schmidt S, Gorski M: LMX1B is essential for the maintenance of differentiated podocytes in adult kidneys. Journal of the American Society of Nephrology 2013, 24(11):1830–1848.

67. Eastman JT, Witmer LM, Ridgely RC, Kuhn KL: Divergence in skeletal mass and bone morphology in antarctic notothenioid fishes. Journal of Morphology 2014, 275(8):841–861.

68. Albertson RC, Yan Y-L, Titus TA, Pisano E, Vacchi M, Yelick PC, Detrich III HW, Postlethwait JH: Molecular pedomorphism underlies craniofacial skeletal evolution in Antarctic notothenioid fishes. BMC evolutionary biology 2010, 10(1):4.

69. Iwami T: Osteology and relationships of the family Channichthyidae. Memoirs of National Institute of Polar Research Ser E, Biology and medical science 1985, 36:1–69.

70. Voskoboinikova O: Evolutionary significance of heterochronies in the development of the bony skeleton in fishes of the suborder Notothenioidei (Perciformes). Journal of Ichthyology 2001, 41(6):415–424.

71. Price PA, Otsuka A, Poser JW, Kristaponis J, Raman N: Characterization of a gamma-carboxyglutamic acid-containing protein from bone. Proceedings of the National Academy of Sciences 1976, 73(5):1447–1451.

72. Tsao Y-T, Huang Y-J, Wu H-H, Liu Y-A, Liu Y-S, Lee OK: Osteocalcin mediates biomineralization during osteogenic maturation in human mesenchymal stromal cells. International journal of molecular sciences 2017, 18(1):159.

73. Debaenst S, Jarayseh T, De Saffel H, Bek JW, Boone M, Josipovic I, Kibleur P, Kwon RY, Coucke PJ, Willaert A: Crispant analysis in zebrafish as a tool for rapid functional screening of disease-causing genes for bone fragility. Elife 2025, 13:RP100060.

74. Eastman JT: The axes of divergence for the evolutionary radiation of notothenioid fishes in Antarctica. Diversity 2024, 16(4):214.

75. Waas WF, Druzina Z, Hanan M, Schimmel P: Role of a tRNA base modification and its precursors in frameshifting in eukaryotes. Journal of Biological Chemistry 2007, 282(36):26026–26034.

76. Fraser KP, Peck LS, Clark MS, Clarke A, Hill SL: Life in the freezer: protein metabolism in Antarctic fish. Royal Society Open Science 2022, 9(3).

77. Takahashi JS: Transcriptional architecture of the mammalian circadian clock. Nature Reviews Genetics 2017, 18(3):164–179.

78. Siepka SM, Yoo S-H, Park J, Song W, Kumar V, Hu Y, Lee C, Takahashi JS: Circadian mutant Overtime reveals F-box protein FBXL3 regulation of cryptochrome and period gene expression. Cell 2007, 129(5):1011–1023.

79. Kim B-M, Amores A, Kang S, Ahn D-H, Kim J-H, Kim I-C, Lee JH, Lee SG, Lee H, Lee J: Antarctic blackfin icefish genome reveals adaptations to extreme environments. Nature ecology & evolution 2019, 3(3):469–478.

80. Lau J, Herzog H: CART in the regulation of appetite and energy homeostasis. Frontiers in neuroscience 2014, 8:313.

81. Rønnestad I, Gomes AS, Murashita K, Angotzi R, Jönsson E, Volkoff H: Appetite-controlling endocrine systems in teleosts. Frontiers in endocrinology 2017, 8:73.

82. Daane JM, Auvinet J, Stoebenau A, Yergeau D, Harris MP, Detrich III HW: Developmental constraint shaped genome evolution and erythrocyte loss in Antarctic fishes following paleoclimate change. PLoS genetics 2020, 16(10):e1009173.

83. Desvignes T, Rivera-Colón AG, Postlethwait JH: Hemoglobin-gene cluster deletions in Antarctic white-blooded icefishes facilitated by transposable elements. Genome Biology and Evolution 2025, 17(10):evaf184.

84. Babst M, Sato TK, Banta LM, Emr SD: Endosomal transport function in yeast requires a novel AAA-type ATPase, Vps4p. The EMBO journal 1997.

85. Di Rosa G, Pustorino G, Spano M, Campion D, Calabrò M, Aguennouz M, Caccamo D, Legallic S, Sgro DL, Bonsignore M: Type I hyperprolinemia and proline dehydrogenase (PRODH) mutations in four Italian children with epilepsy and mental retardation. Psychiatric genetics 2008, 18(1):40–42.

86. Hatano M, Udagawa T, Kanamori T, Sutani A, Mori T, Sohara E, Uchida S, Morio T, Nishioka M: A novel SLC5A2 heterozygous variant in a family with familial renal glucosuria. Human Genome Variation 2022, 9(1):42.

87. Allaire P, Fox J, Kitchner T, Gabor R, Folz C, Bettadahalli S, Hebbring S: Familial renal glucosuria and potential pharmacogenetic impact on Sodium-Glucose Cotransporter-2 inhibitors. Kidney360 2025, 6(4):521–530.

88. Zhang L, Hirano A, Hsu P-K, Jones CR, Sakai N, Okuro M, McMahon T, Yamazaki M, Xu Y, Saigoh N: A PERIOD3 variant causes a circadian phenotype and is associated with a seasonal mood trait. Proceedings of the National Academy of Sciences 2016, 113(11):E1536–E1544.

89. Plavc L, Skubic C, Dolenc Grošelj L, Rozman D: Variants in the circadian clock genes PER2 and PER3 associate with familial sleep phase disorders. Chronobiology international 2024, 41(5):757–766.

90. Seu KG, Trump LR, Emberesh S, Lorsbach RB, Johnson C, Meznarich J, Underhill HR, Chou ST, Sakthivel H, Nassar NN: VPS4A mutations in humans cause syndromic congenital dyserythropoietic anemia due to cytokinesis and trafficking defects. The American Journal of Human Genetics 2020, 107(6):1149–1156.

91. Shipman A, Gao Y, Liu D, Sun S, Zang J, Sun P, Syed Z, Bhagavathi A, Smith E, Erickson T: Defects in exosome biogenesis are associated with sensorimotor defects in zebrafish vps4a mutants. Journal of Neuroscience 2024, 44(50).

92. Holeton GF: Oxygen uptake and circulation by a hemoglobinless Antarctic fish (Chaenocephalus aceratus lonnberg) compared with three red-blooded Antartic fish. Comparative biochemistry and physiology 1970, 34(2):457–471.

