## Supplementary_Figures for "Gene loss under constant cold reveals “natural knockout” loci in Antarctic notothenioid fishes": Supplementary_Figures.docx

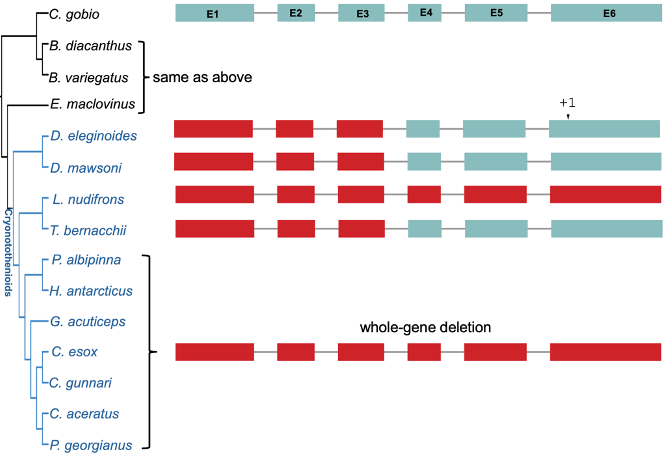

**Supplementary Figure 1.** Schematic representation of the *tyw3* gene loss showing multiple exon deletion across Antarctic notothenioids

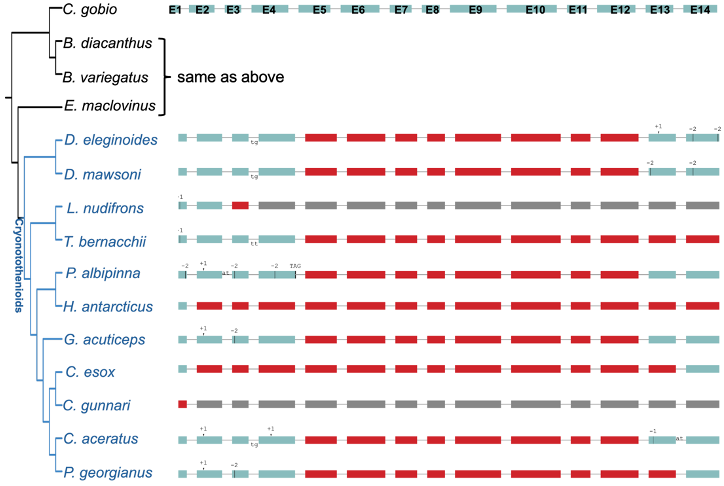

**Supplementary Figure 2.** Schematic representation of the *tyw4* gene loss showing multiple exon deletion and frameshift mutations across Antarctic notothenioids

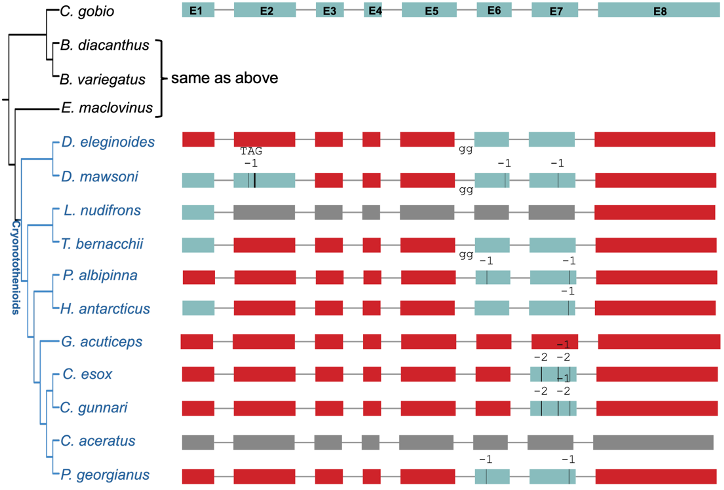

**Supplementary Figure 3.** Schematic representation of the *tyw5* gene loss showing multiple exon deletion across Antarctic notothenioids

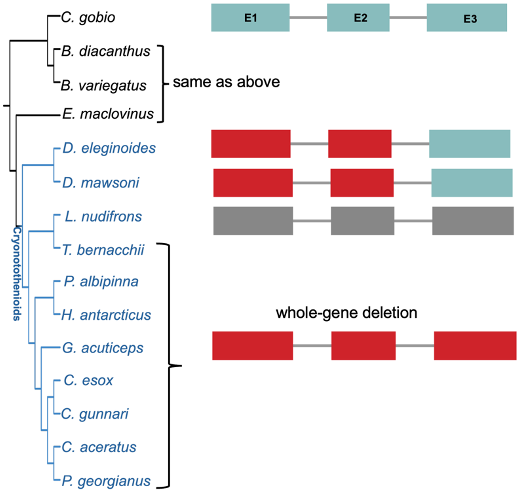

**Supplementary Figure 4.** Schematic representation of the *cart2* gene loss showing multiple exon deletion across Antarctic notothenioids

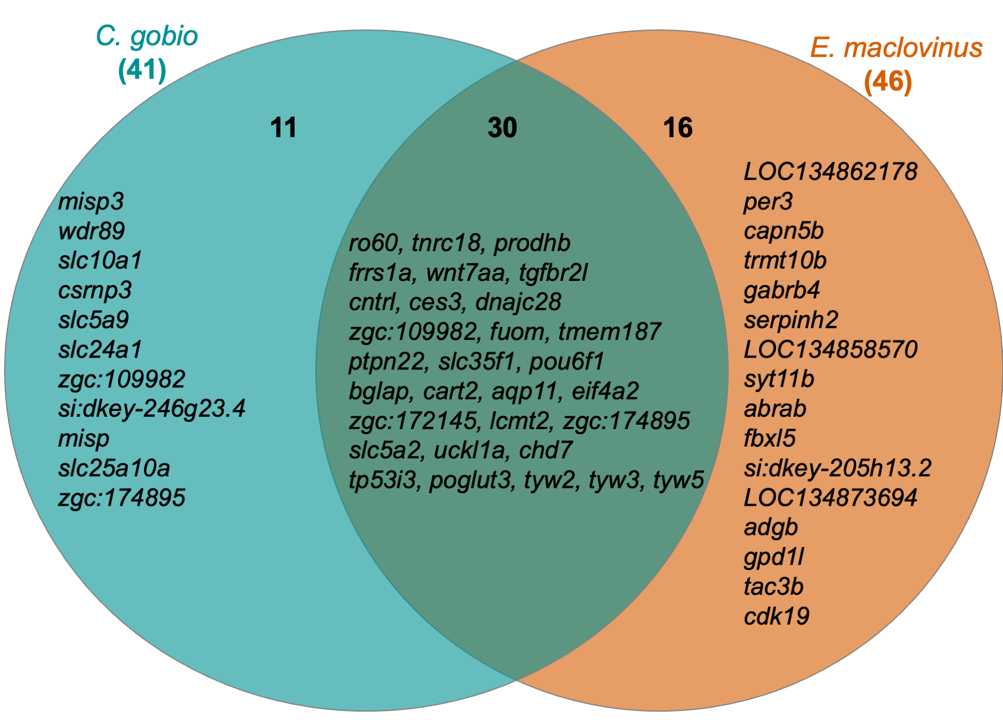

**Supplementary Figure 5.** Venn diagram showing the predicted inactivated single copy orthologs genes in Antarctic clade using both references.

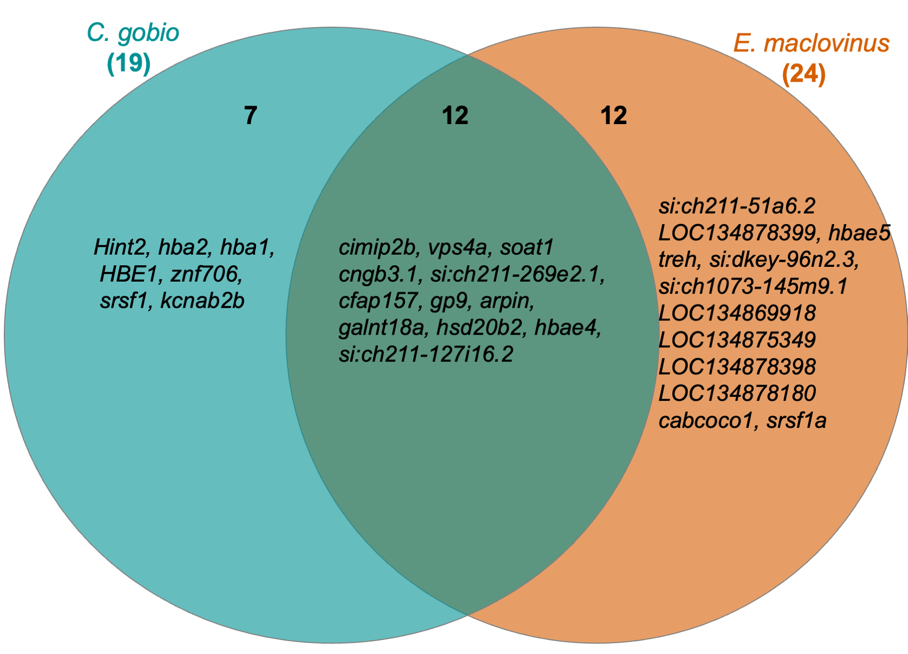

**Supplementary Figure 6.** Venn diagram showing the predicted inactivated single copy orthologs genes in icefish family using both references.
